# EnsAgent: a tool-ensemble multiple Agent system for robust annotation in spatial transcriptomics

**DOI:** 10.64898/2026.03.10.710824

**Authors:** Dongjian Zhang, Mingshen Zhang, Nuo Li, Chaoxin Zheng, Lixin Liang, Xiao Ke, Qishi Dong

## Abstract

**Motivation:** Automated domain annotation in spatially resolved transcriptomics (SRT) remains challenging since it depends on gene expression, morphology, and clinical conventions, which vary across cohorts and platforms. While Large Language Model (LLM)-driven agents show promise, current approaches typically condition semantic reasoning on static, single-method partitions. This reliance makes annotation pipelines fragile to upstream partition errors and prone to hallucinations when molecular evidence is ambiguous. A robust framework integrating ensemble intelligence with iterative, evidence-based reasoning is required to ensure reproducibility and accuracy.

**Results:** We introduce EnsAgent, a tool-ensemble multi-agent system designed for robust SRT annotation. Uniquely, EnsAgent decouples structural partitioning from semantic labeling via a Consultation–Review workflow. A Tool-Runner Agent orchestrates a diverse portfolio of clustering algorithms via the Model Context Protocol (MCP), generating a consensus partition optimized by a multimodal Scoring Agent. Subsequently, a Proposer–Critic feedback loop coordinates four specialized experts (Marker, Pathway, Spatiality, and Visual) to formulate annotations with explicit evidence trails and uncertainty estimates. Benchmarking on three SRT datasets demonstrates that EnsAgent effectively neutralizes batch effects and resolves subtle tumor microenvironment niches missed by single-paradigm baselines, delivering state-of-the-art accuracy and interpretability.

**Availability and Implementation:** EnsAgent is available at github.com/keviccz/ensAgent.

**Supplementary information:** Supplementary data are available at Bioinformatics online.

## 1 Introduction

The rapid emergence of Spatially Resolved Transcriptomics (SRT), such as 10x Visium [Ståhl et al., 2016], Slide-seq [Rodriques et al., 2019], Stereo-seq [Chen et al., 2022], and MERFISH [Chen et al., 2015], enables a spatially grounded view of biological organization, revealing how cells assemble into higher-order structures and how tissue architecture changes across development [Moses and Pachter, 2022, Bressan et al., 2023]. To translate these rich measurements into biological insight, a crucial step is domain annotation, which assigns biologically meaningful labels, such as anatomical compartments, pathological lesions, or microenvironmental states, to spatially coherent regions, thereby providing a semantic layer that bridges computational outputs and biological knowledge. However, biologically meaningful domains are defined not only by gene expression signatures but also by morphological context and clinical conventions that vary by organ and disease subtype, leading to inconsistent annotations across cohorts and platforms. This creates a steep knowledge barrier and frequently demands strong clinical and pathological expertise.

To alleviate the expertise barrier, there has been a growing interest in leveraging multimodal large language models (MLLMs) for domain annotation. Existing studies largely follow three recurring paradigms. First, tool-augmented agents, best exemplified by STAgent [Lin et al., 2025] and SpatialAgent [Wang et al., 2025], use MLLMs to orchestrate established spatial domain detection pipelines and summarize the outputs. Second, representation-based prompting compresses high-dimensional spatial omics into language-compatible inputs. For example, OmicsNavigator [Li et al., 2025] encodes regions of interest (ROIs) as textual descriptors for Large Language Model (LLM)-based interpretation, while LLMiniST [Wei et al., 2025] leverages local microenvironment cues and cell-type composition via zero-shot prompting to identify niches. While these systems can improve interpretability using MLLMs, they typically condition annotation on a single upstream partitioning or assignment [Wang et al., 2025, Lin et al., 2025, Li et al., 2025, Wei et al., 2025] and then provide the MLLM with compressed evidence to produce semantic annotations. Such reliance on single-method upstream partitions makes the annotation pipeline fragile to resolution mismatch, batch effects across slides, tissue-specific heterogeneity, and hyperparameter sensitivity across SRT data. When evidence is incomplete or conflicting, MLLM-based annotators may overfit to superficial marker cues, produce unstable labels across datasets, or exhibit hallucinations.

In addition, many agentic frameworks treat semantic labeling as a definitive, one-shot inference, compressing molecular evidence and contextual cues into a single generation step lacking explicit arbitration to reconcile data-driven signals with potential hallucinations [Dip and Zhang, 2025, Wei et al., 2025, Li et al., 2025, Wang et al., 2025]. These systems typically return narrative labels without a structured evidence trail or decision logs, limiting reproducibility and auditability. Moreover, current systems generally output a single label set rather than calibrated probabilities that quantify uncertainty. They typically do not explicitly provide confidence measures that account for the stability of the underlying spatial partition itself. Consequently, without granular feedback loops to flag low-fidelity annotations, errors propagate downstream, compromising region-specific marker discovery and cross-sample harmonization.

In order to address these limitations, we introduce **EnsAgent**, a tool-ensemble multiple agent system for robust annotation in spatial transcriptomics. The system follows a Consultation–Review workflow spanning three stages. In Stage 1, a Tool-Runner Agent executes a portfolio of spatial domain detection tools, each encapsulated as a callable tool via the Model Context Protocol (MCP), producing a diverse set of candidate partitions. In Stage 2, a Scoring Agent evaluates each candidate using multimodal evidence to construct a score-weighted ensemble, yielding a fine-grained consensus domain map. In Stage 3, a Proposer Agent coordinates four specialized experts to formulate candidate labels with evidence-linked uncertainty estimates. A Critic Agent subsequently audits these claims against a knowledge base (KB), identifying inconsistencies and triggering targeted re-analysis by specific experts. By coupling ensemble-based structural robustness with iterative, evidence-grounded multi-agent reasoning, EnsAgent delivers accurate annotations resilient to platform heterogeneity. We demonstrate this superiority in diverse contexts, including recovering the fine-grained laminar architecture and functional logic of the human cortex [Maynard et al., 2021], capturing subtle tumor microenvironment subgroups in human breast cancer sections [Ståhl et al., 2016] missed by single-paradigm baselines, and neutralizing batch effects in Mouse Olfactory Bulb tissues [Chen et al., 2022].

## 2 Methods

### 2.1 Overview

EnsAgent is a role-specialized multi-agent system for robust spatial domain annotation that couples multi-method ensembling with iterative evidence verification. It takes SRT inputs provided as an AnnData object, together with a pre-configured analysis environment and a natural-language context prompt specifying key biological metadata (e.g., species, tissue, platform, and disease condition). A Tool-Runner Agent interfaces with an MCP-based tool layer to execute a portfolio of domain discovery methods and downstream analyses within a reproducible coding backend. Their outputs are then integrated by a Scoring Agent via a dual-stream scoring engine that combines quantitative spatial statistics with visual and morphological assessments to construct a score-weighted consensus domain map. For semantic annotation, a Proposer Agent coordinates specialized experts to generate top-*k* label hypotheses with evidence-linked uncertainty estimates, while a Critic Agent audits these claims against curated knowledge and cross-evidence consistency, triggering targeted re-analysis when evidence is insufficient or contradictory. The Consultation–Review loop iterates until convergence, returning an annotated domain map along with structured JSON outputs that retain scores, evidence, and decision traces for auditability. Overall runtime is dominated by backend tool execution, with an additional contribution from Vision Language Model (VLM)-based image evaluation. In contrast, semantic annotation scales with the length of the LLM dialogue rather than the number of spots, contributing *<* 5% overhead(Supplementary Note S1.10). Detailed prompts, methodological specifications, and complexity analyses are provided in Supplementary Note S1.

### 2.2 Tool-Runner Agent: MCP-based Tool Orchestration and Standardized Candidate Partitions

The Tool-Runner Agent converts SRT inputs into a standardized collection of candidate domain partitions and evidence artifacts for downstream scoring and annotation. Let 𝒟 denote the input AnnData object (with *n* spots), *c* a natural-language context prompt (Supplementary Note S1.11.1) specifying biological metadata, and *E* the configured execution environment. The Tool-Runner Agent maintains a tool library *𝒯* = {*𝒯* _1_, …, *𝒯*_*M*_ *}*, where each tool {_*m*_ encapsulates a spatial domain discovery method behind a MCP interface, including BayesSpace [Zhao et al., 2021], BASS [Li and Zhou, 2022], DR-SC [Yu et al., 2022], STAGATE [Dong and Zhang, 2022], stLearn [Pham et al., 2023], GraphST [Long et al., 2023], IRIS [Ma and Zhou, 2024], and SEDR [Xu et al., 2024]. Each tool defines a callable map 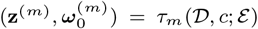, where **z**^(*m*),^ is an *n*-length spot-level domain assignment with *K*_*m*_ clusters and 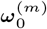 contains tool-native provenance. Given (*𝒟, c*), the Tool-Runner Agent SRT constructs an execution plan by selecting a portfolio ℳ (*c*) *⊆ {*1, …, *M}* (default: a fixed portfolio; optionally conditioned on platform and tissue keywords in *c*) (Supplementary Note S1.5.1). It dispatches MCP calls to all selected tools and collects their outputs: 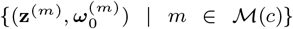, thereby producing heterogeneous candidate partitions with standardized I/O.

Because unsupervised methods assign arbitrary label indices (e.g., “Cluster 1” in one method may correspond to “Cluster 5” in another), 𝒜_TR_ aligns each candidate labeling **z**^(*m*),^ into a shared label space defined by a reference partition **r** (Supplementary Note S1.5.2). Instead of heuristic matching, we formulate this as a maximum-weight assignment problem based on the Intersection-over-Union (IoU) metric, solved globally via the Hungarian algorithm [Kuhn, 1955]. To ensure the alignment step is reproducible and backend-agnostic, Tool-Runner Agent encapsulates this routine as an MCP-callable utility tool, denoted *τ*_align_, producing the aligned label 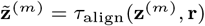.

Subsequently, it invokes additional MCP-wrapped analysis tools to generate a common set of lightweight evidence artifacts required by downstream agents. These include: (i) spatial domain visualizations for each candidate partition; (ii) domain-level marker statistics based on differentially expressed genes (DEGs); and (iii) functional enrichment profiles based on GO [Ashburner et al., 2000] and KEGG [Kanehisa and Goto, 2000]. All artifacts are serialized as CSV/JSON tables and PNG figures with tool identifiers, configuration hashes, and provenance metadata, and are appended to the per-method provenance bundle: 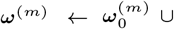 *{*alignment metadata, evidence artifacts*}*. Eventually, the Tool-Runner Agent returns a manifest:

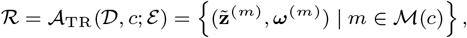

Where 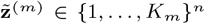 denotes the aligned spot-level domain assignment for method *m*, and ***ω***^(*m*),^ is the corresponding provenance bundle, including (i) alignment metadata, (ii) evidence artifacts (plots, DEGs, enrichment tables), and (iii) execution provenance such as runtime metadata, intermediate embeddings, and diagnostic logs (Supplementary Note S1.12.1).

### 2.3 Scoring Agent:Dual-Stream Quality Evaluation and Consensus Aggregation

The Scoring Agent *A*_SC_ functions as an integration kernel, mapping the set of aligned candidate partitions *ℛ* produced by the Tool-Runner into a single consensus domain map **y**^*∗*,^ through a reliability-weighted voting mechanism. It comprises an Evaluation Module ℳ_eva_, which queries an MLLM evaluator to distill molecular and spatial evidence into normalized quality scores, and a Visual Module ℳ _vis_, which queries a vision-capable MLLM to assess morphological fidelity (prompt provided in Supplementary Note S1.11.2). Let ω ^(*m,k*),^ denote the domain-*k* evidence bundle extracted from *ω* ^(*m*),^. For each candidate partition *m* and its domain *k ∈ {*1, …, *K*_*m*_*}*, the Evaluation Module maps the provenance bundle *ω* ^(*m*),^ to a compact evidence descriptor:

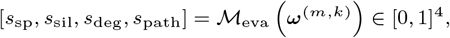

where the four normalized domain-quality features quantify spatial coherence (*s*_sp_), cluster separability (*s*_sil_), marker specificity (*s*_deg_), and pathway consistency (*s*_path_). Simultaneously, Visual Module ℳ_vis_ incorporates morphological reasoning while preventing visual priors from overriding molecular signals. Let **I**_*m,k*_ *∈ ω* ^(*m*),^ denote the visualized spatial map corresponding to domain *k*. The Visual Module outputs three morphology-related factors:

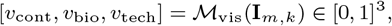

capturing geometric continuity, biological plausibility, and technical artifact likelihood, respectively. The Scoring Agent forms a baseline reliability score as a linear combination of the four evaluation features:

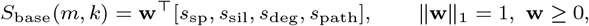

and summarize the visual factors into a bounded modulation term *v*(*m, k*) via a linear aggregation of [*v*_cont_, *v*_bio_, *v*_tech_] followed by rescaling (Supplementary Note S1.6.1). The final domain reliability score is then obtained by conservatively modulating the baseline score with the visual term:

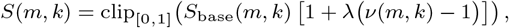

where *λ* controls the sensitivity to visual modulation.

The consensus map **y**^*∗*,^ *∈ {*1, …, *K*^*∗*,^*}*^*n*^ is constructed via spot-wise weighted voting over the aligned candidate partitions. For each spot *u* 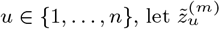 denote the aligned label assigned by method *m* ∈*ℳ*_(*c*)_. We define a dynamic voting weight by the reliability of the predicted domain 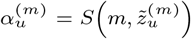, thereby methods contribute more strongly in regions where their corresponding domains are scored as reliable. The consensus label is then given by

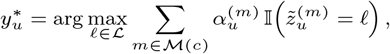

where ℒ = 𝒰_*m∈* ℳ (*c*)_^*{*1, …, *Km}*,^ is the union of aligned label indices.

To quantify local stability, we compute a spot-wise disagreement score:

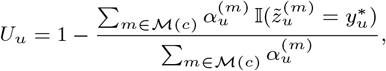

where a higher *U*_*u*_ indicates stronger cross-method conflict, typically concentrated near ambiguous boundaries. A spatial smoothing operator ℱ_knn_ (using majority voting within the 20-nearest neighbors) is applied to **y**^*∗*,^ to suppress isolated stochastic assignments while preserving contiguous domain structure (Supplementary Note S1.6.4). Finally, we re-invoke an MCP-wrapped analysis toolchain *τ*_analy_ on the consensus labels to generate standardized artifacts for downstream annotation:

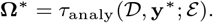

The resulting bundle **Ω**^*∗*,^ contains updated domain-level DEGs, enrichment profiles, and spatial/morphology visualizations specific to the consensus partition, and serves as the shared evidence substrate for the subsequent Proposer–Critic annotation stage.

### 2.4 Multi-expert evaluation and Critic–Proposer iterative refinement loop

We implement a semantic interpretation framework that couples multi-expert probabilistic evaluation with a Critic–Proposer iterative refinement loop. The Proposer aggregates evidence to generate the strongest label hypotheses, whereas the Critic performs robustness-oriented auditing that searches for cross-modality contradictions and requests targeted re-analysis when support is insufficient (Supplementary Note S1.11.3). For each consensus domain *k*, let Ω_*k*_ *∈* **Ω**^*∗*,^ denote the domain-restricted evidence bundle, and let 𝒴 denote the candidate anatomical label set. The Proposer Agent coordinates a set of experts *𝒢*=*{*Marker, Pathway, Spatiality, Visual}, where each expert *e ∈ 𝒢* queries an MLLM judge on its modality-specific evidence in Ω_*k*_ and outputs a calibrated posterior probability over labels:

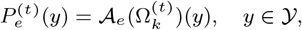

where *t* indexes refinement iterations, with 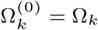, and 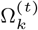 may be updated by Critic-triggered re-analysis. The Proposer synthesizes expert posteriors via a linear opinion pool:

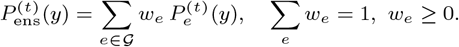

It proposes *top-K* label candidates 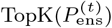and outputs the primary hypothesis:

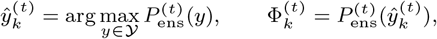

where 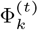 denotes the ensemble confidence for domain *k* at iteration *t*.

To make the decision robust to modality-specific failure modes, the Critic evaluates the Proposer’s hypothesis using a worst-case support score and an explicit cross-modality consistency check. Define the worst-case support among experts as:

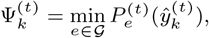

thereby a label cannot pass if any modality assigns it a low probability. In addition, the Critic computes a validation score 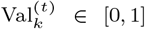 to strictly quantify the objective validity of the annotation by integrating inter-expert consensus with external biological verification:

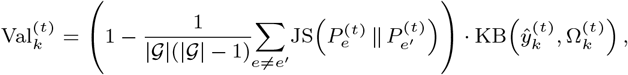

where JS(*·∥·*) is Jensen-Shannon divergence representing expert consistency. Knowledge Base quantifies the biological plausibility of the proposed label, denoted as KB(*·*) *∈* [0, 1]. This adaptive repository is dynamically synthesized by the LLM to encode dataset-specific logic and supports manual curation, employing a synchronous update mechanism to rigorously enforce biological constraints (Supplementary Note S1.7.3).

Finally, the Critic forms an audit score that combines Proposer confidence with robustness checks:

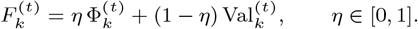

If 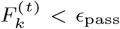 the Critic triggers targeted refinement by selecting the most conflicting expert:

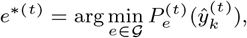

and issuing an action 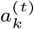 (a natural-language instruction plus tool calls) to request additional domain-specific analyses, such as re-run DEGs with stricter thresholds, add alternative pathway collections, compute spatial autocorrelation, or inspect histology overlays. This updates the evidence bundle 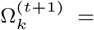 Update 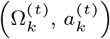, after which the Proposer recomputes 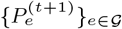 and the Critic re-audits. The loop terminates when 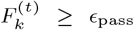the top label 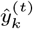 stabilizes, or a maximum iteration budget *t* = *T*_max_ is reached (Supplementary Note S1.7.4). Upon termination, the system emits a structured annotation manifest 𝒪 _final_. For each domain *k*, 𝒪 _final_(*k*) contains the validated label *ŷ*_*k*_, the audit score *F*_*k*_, the top-*K* candidates *C*_*k*_, modality-specific posteriors *{P*_*e*_(*y*)*}*_*e∈𝒢*_, and an evidence-linked decision trace (including Critic flags and any triggered re-analysis), enabling reproducibility, auditability, and error localization, with detailed output examples provided in Supplementary Note S1.12.

## 3 Results

### 3.1 Accurate Lamination and Functional Characterization in the Human Cortex

To evaluate the accuracy of EnsAgent in delineating cortical laminar organization, we applied it to the human dorsolateral prefrontal cortex (DLPFC) 10x Visium dataset [Maynard et al., 2021]. Uniquely, the framework orchestrates a Tool-Runner Agent to generate standardized candidate partitions from established algorithms, followed by a dual-stream Scoring Agent that acts as an autonomous referee (Fig. 2a). By modulating a quantitative biological baseline (Base Score, *S*_base_) with a vision-language model-derived morphological factor (*v*), the system downweights incoherent partitions to construct a robust consensus domain map. This reliability-weighted aggregation strategy ensures anatomical fidelity by penalizing the fragmentation often missed by unimodal metrics.

**Figure 1.**
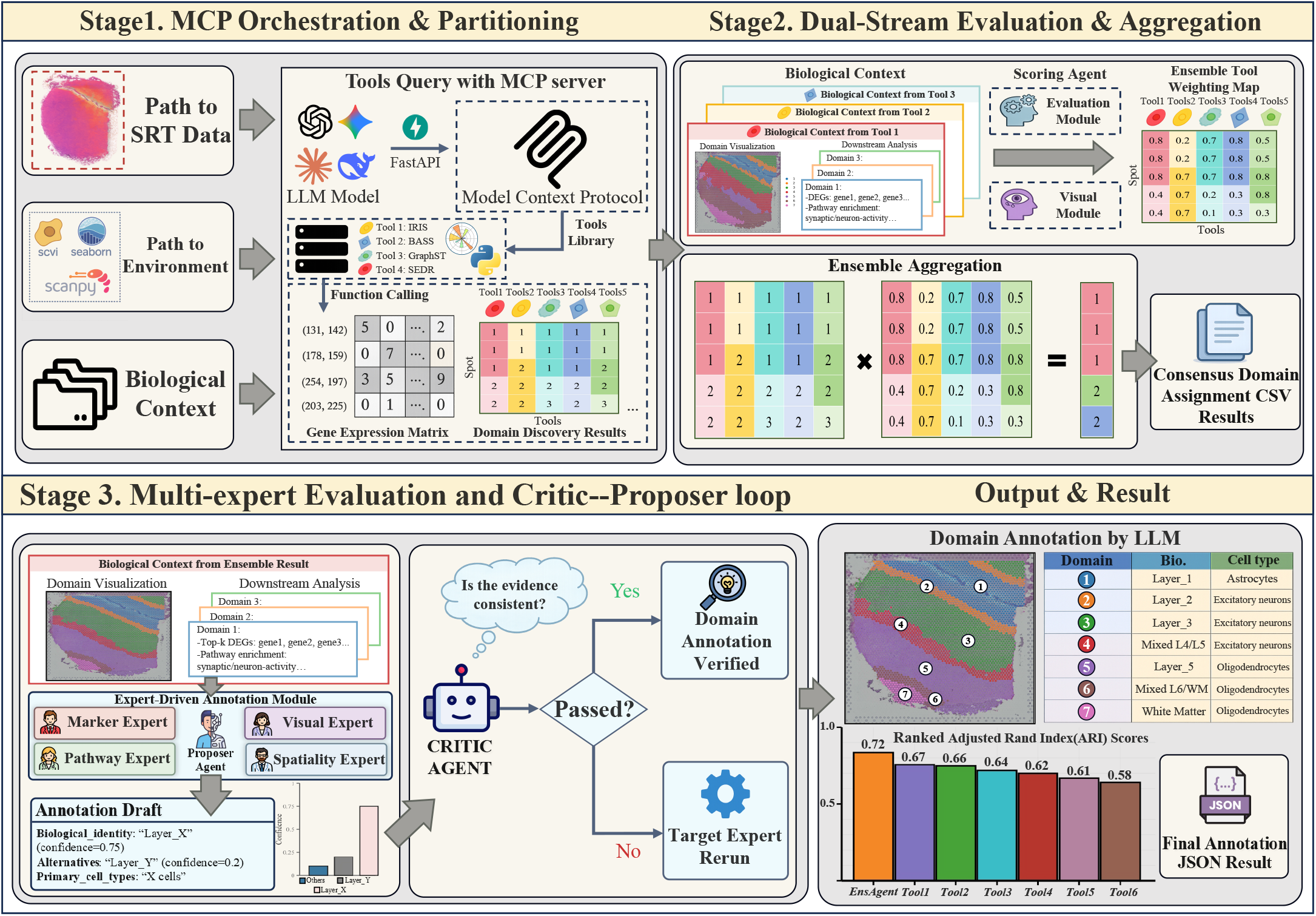
Overview of the EnsAgent Framework. **Stage 1: Tool-Runner Agent: MCP-based Tool Orchestration and Standardized Candidate Partitions**. The MCP-based module orchestrates diverse tools (e.g., IRIS, BASS) to generate standardized candidate partitions from raw SRT data. **Stage 2: Scoring Agent: Dual-Stream Quality Evaluation and Consensus Aggregation**. A dual-stream engine integrates biological fidelity (*S*_base_) and visual constraints to aggregate weighted predictions into a robust consensus. **Stage 3: Multi-expert evaluation and Critic–Proposer iterative refinement loop**. A Proposer–Critic loop coordinates four experts (Marker, Visual, Pathway, Spatiality) to refine evidence and produce verified domain identities. **Alt Text:** Flowchart showing EnsAgent’s three stages: (1) Tool-Runner Agent generating partitions; (2) Scoring Agent calculating consensus; (3) Proposer-Critic loop for final annotation.

**Figure 2.**
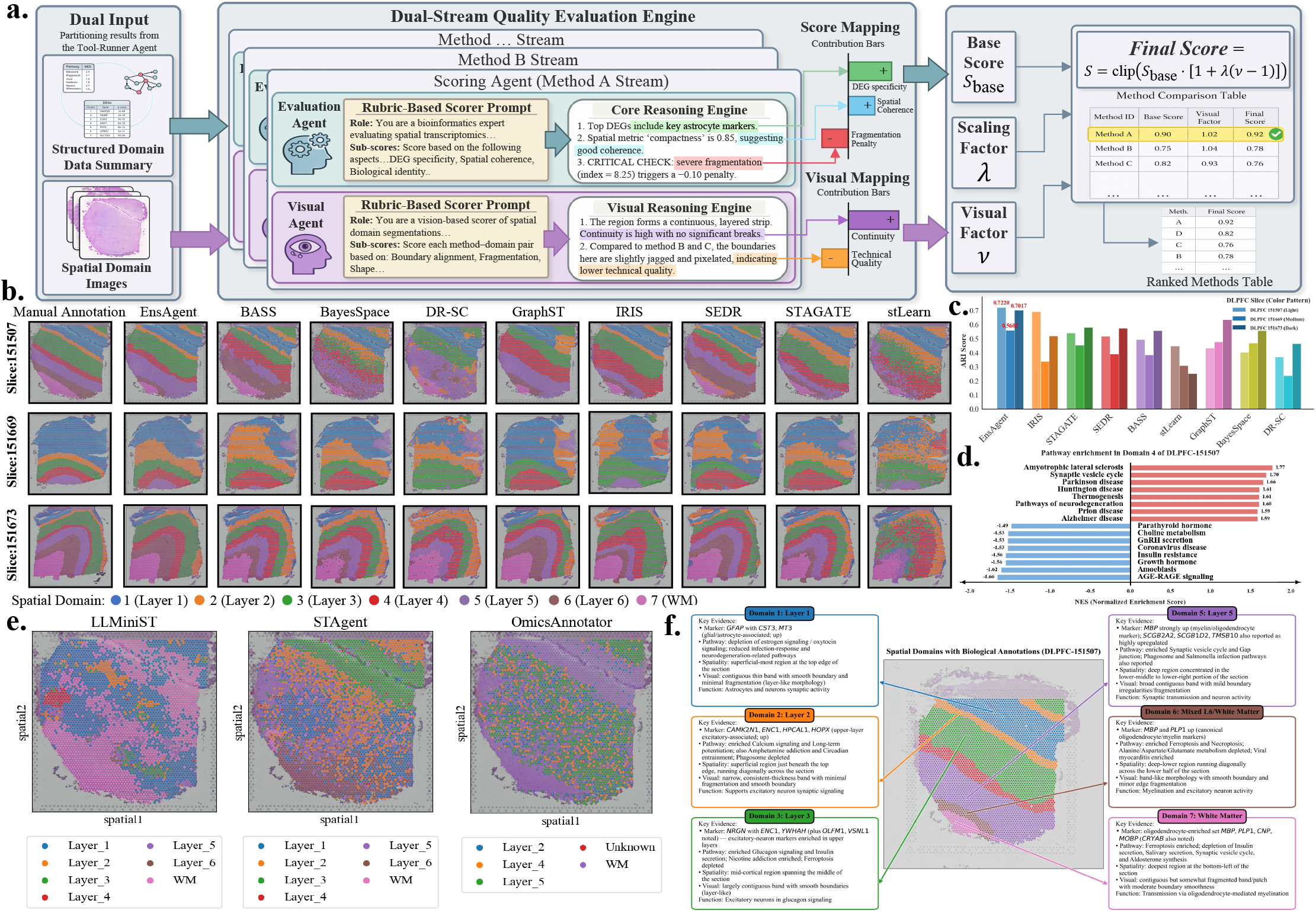
**(a)** Schematic of the Dual-Stream Quality Evaluation Engine, which selects optimal partitions by synthesizing biological fidelity (Base Score, *S*_base_) and visual morphological constraints (Visual Factor, *v*). The final score is computed by modulating the base score with the visual factor using a scaling parameter *λ*. **(b)** Spatial domain visualization of EnsAgent vs. manual annotations and 8 baselines across DLPFC slices 151507, 151669, and 151673. **(c)** Quantitative benchmarking (ARI/NMI) demonstrating EnsAgent’s consistent improvement. **(d)** Pathway enrichment for Domain 4 identifying neuronal programs. **(e)** Visual comparison against LLM-driven agents (OmicsAnnotator [Li et al., 2025], STAgent [Wang et al., 2025], LLMiniST [Wei et al., 2025]). **(f)** Example of evidence-based annotation output, where the Four-Expert Module provides molecular and functional reasoning for domain identity. **Alt Text:** Composite figure: (a) scoring schematic; (b) domain maps comparison; (c) ARI benchmarks; (d) pathway bar charts; (e) agent comparison maps; (f) expert reasoning text.

Visually, baselines such as DR-SC [Yu et al., 2022] and stLearn [Pham et al., 2023] exhibited noisy, fragmented partitions across slices 151507, 151669, and 151673, leading to layer mixing. BayesSpace [Zhao et al., 2021] and IRIS [Ma and Zhou, 2024] showed incomplete separation for slice 151669. In contrast, EnsAgent recovers spatially coherent layers with sharp boundaries, closely matching manual annotations (Fig. 2b). Quantitatively, EnsAgent achieves the highest Normalized Mutual Information (NMI) with a tight distribution, indicating strong cross-slice robustness (Supplementary Fig. S1). Furthermore, Adjusted Rand Index (ARI) comparisons consistently rank EnsAgent as the top-performing method (mean ARI = 0.67), showing an 4.31%–17.39% improvement over individual baselines (Fig. 2c), confirming its robustness to anatomical variability and technical perturbations.

Beyond metrics, the Proposer Agent provides biological evidence supporting domain annotation. For Domain 4 on the DLPFC slice 151507 (Fig. 2d), it identifies enrichment dominated by neuronal functions such as “Synaptic vesicle cycle” (NES = 1.70) as well as neurodegeneration-related pathways, reflecting shared mitochondrial and synaptic activity modules prevalent in high-activity neurons. This aligns with metabolically demanding cortical neurons, providing mechanistic support for the laminar identity. Conversely, the negative enrichment of non-neural signals (e.g., “Parathyroid hormone”) confirms the agent prioritizes domain-relevant programs over off-target systemic noise.

We further benchmarked structural fidelity against recent bioinformatics agents (Fig. 2e). Visual benchmarking reveals that baseline agents struggle to balance coherence with specificity. OmicsAnnotator [Li et al., 2025] collapsed the original seven layers into five, leaving significant regions as uninformative “Unknown” clusters due to a lack of spatial constraints. Similarly, STAgent [Wang et al., 2025] exhibited reduced spatial fidelity, notably failing to resolve the boundaries between superficial layers (L1–L3), resulting in a noisy, intermixed distribution that disrupted the continuous laminar progression. Finally, while LLMiniST [Wei et al., 2025] captured broader identities, it suffered from severe “salt-and-pepper” fragmentation, failing to reconstruct band-like morphologies.

In contrast, EnsAgent employs a distinct Dual-Agent Scoring Engine that integrates biological marker evaluation with MLLM-driven visual constraints to penalize fragmentation. By effectively filtering out low-quality partitions, our method achieves segmentation that demonstrates superior alignment with the ground truth. The Four-Expert Annotation Module generates biologically transparent definitions (Fig. 2f), replacing the simple class labels typical of baselines with comprehensive “Key Evidence” synthesized from experts. For Domain 3, the system cited the upregulation of *NRGN* and *ENC1* alongside “Glucagon signaling” pathways to definitively identify “Layer 3”. Furthermore, the system successfully resolved the deep-layer complexity that confused STAgent; it distinguished Domain 6 (“Mixed L6/White Matter”), which is characterized by mixed myelination and excitatory neuron activity, from the pure Domain 7 (“White Matter”), defined strictly by oligodendrocyte-mediated myelination markers (e.g., *MBP, PLP1, CNP* ). This demonstrates that integrating EnsAgent can capture the full morphological semantics of the cortex.

To assess robustness and the contribution of individual modules, we performed extensive ablation and sensitivity analyses (Supplementary Table S2). Running the full framework 10 times yielded consistently high performance (ARI = 0.6998–0.7220). In contrast, removing the visual module led to a clear drop in accuracy (ARI = 0.6718–0.7048), underscoring the contribution of morphological cues to boundary refinement. Sensitivity analyses further confirmed robustness, with ARI remaining stable (0.6788–0.7176) across a range of visual weights and LLM temperatures. Together, these results indicate that EnsAgent achieves state-of-the-art accuracy while remaining resilient to hyperparameter variation.

### 3.2 Deciphering Tumor Microenvironment Heterogeneity and Signaling in Human Breast Cancer

In addition, we applied EnsAgent to a human breast cancer 10x Visium dataset [Ståhl et al., 2016] and identified 20 distinct spatial domains, refining broad pathologist annotations (Fig. 3a) into more specific functional niches (Fig. 3b). Beyond coarse regional labels, EnsAgent further resolved the immune landscape into eight Immune-Enriched Regions (IERs) and delineated subtle Tumor–Stroma Interfaces (TSIs) alongside deep tumor cores. This high-resolution partitioning achieved a leading ARI (0.6770) and highly competitive NMI (0.7152) relative to manual annotations (Fig. 3c), significantly outperforming individual baseline methods in overall structural coherence (Supplementary Fig. S2). This suggests that EnsAgent effectively integrates complementary domain hypotheses while down-weighting unstable partitions.

**Figure 3.**
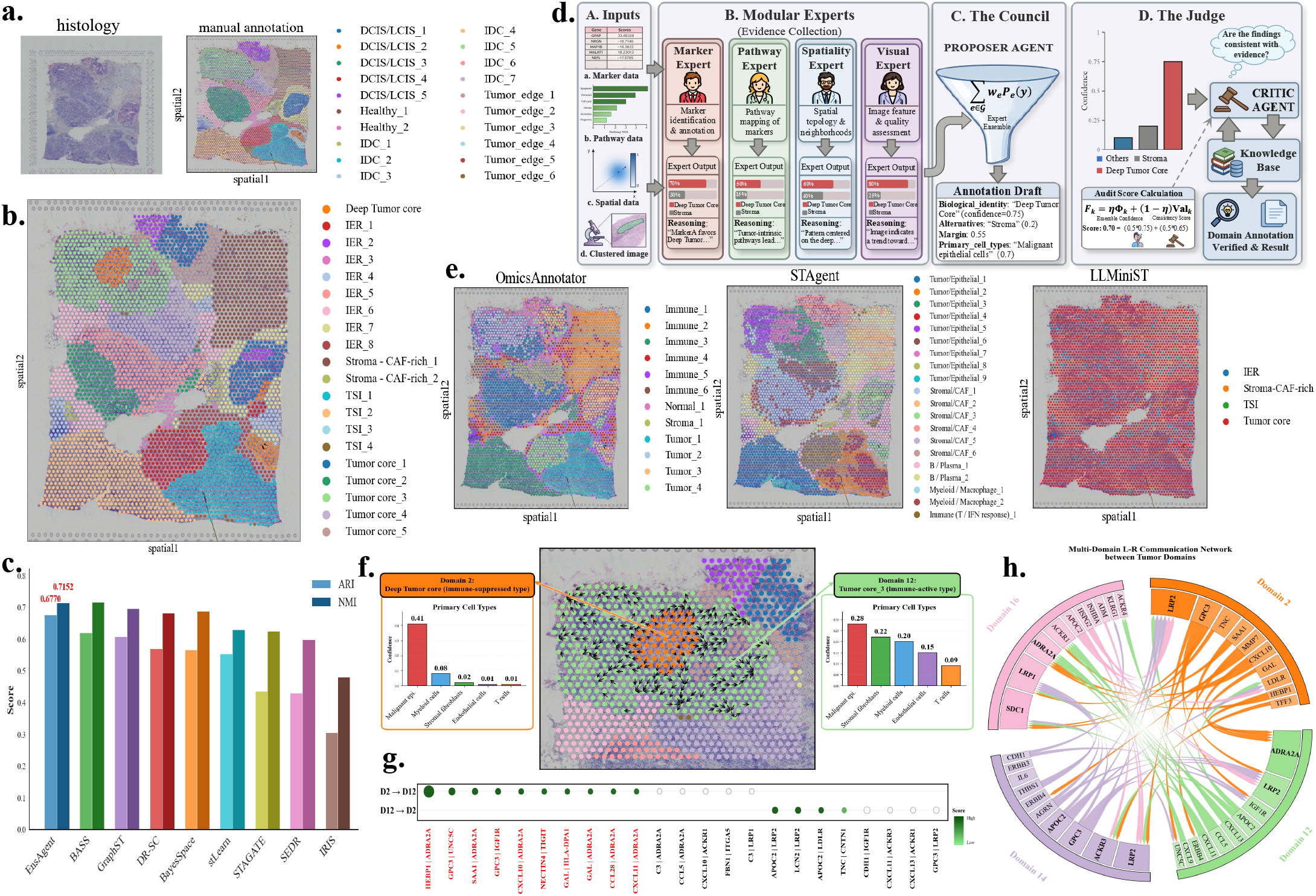
**(a)** H&E staining with pathologist annotations. **(b)** EnsAgent stratification revealing 20 fine-grained spatial domains. **(c)** Benchmarking against manual annotations (ARI/NMI). **(d)** Schematic of the Expert-Proposer-Critic collaborative framework for spatial domain annotation. **(e)** Semantic resolution comparison vs. other agents. **(f)** Spatial vector field illustrating ligand–receptor communication directionality between the Deep Tumor Core (Dom 2) and Tumor–Stroma Interface (Dom 12), with bar charts show distinct cellular compositions for two domains. **(g)** Dot plot ranking significant L–R interactions. **(h)** Chord diagram summarizing domain-specific signaling modules. **Alt Text:** (a) H&E histology and manual annotations; (b) EnsAgent domains; (c) metric boxplots; (d) workflow diagram; (e) agent comparisons; (f) communication vector field; (g) interaction dot plot; (h) signaling chord diagram.

To validate biological insight (Fig. 3d), we benchmarked against the same set of baseline agents (Fig. 3e). OmicsAnnotator produced an oversimplified taxonomy, masking critical transition zones, while STAgent suffered from “semantic redundancy,” generating uninterpretable numerical lists (e.g., “Tumor 1–9”). Notably, LLMiniST exhibited severe “semantic collapse,” erroneously classifying the entire section as “Tumor Core” and obliterating the distinction between necrotic cores and invasive margins.

Uniquely, EnsAgent dissects functionally distinct subregions often conflated by single-method partitions. For instance, while pathologist annotations group Domains 2 and 12 into a single tumor compartment, the Proposer Agent reveals marked differences in cellular composition (Fig. 3f). Domain 2 appears as a malignant-dominant “immune-suppressed” deep core (malignant epi. confidence 0.41), whereas Domain 12 is identified as an “immune-active” niche. This distinction is corroborated by the notable enrichment of T cells in Domain 12 (confidence 0.09) compared to their depletion in Domain 2 (0.01), alongside balanced contributions from stromal fibroblasts (0.22) and myeloid cells (0.20), indicating a tumor–stroma transition. We visualized the underlying molecular logic via spatial vector fields and ranked interaction dot plots (Fig. 3g). The analysis reveals a signaling cascade from Domain 2 to 12, driven by oncogenic pairs (*GPC3* –*UNC5C* ) and an adrenergic module (*CXCL10/11* –*ADRA2A*). Crucially, the agent prioritized immune-regulatory signals like *NECTIN4* –*TIGIT* and *GAL*–*HLA-DPA1*, validating Domain 12 as an immune-recruiting niche [Tekpli et al., 2019]. Finally, the cross-domain L-R network (Fig. 3h) summarizes global signaling patterns, showing that Domain 2 exhibits a *MIF/FN1* -dominated outgoing profile consistent with remodeling activity, while Domains 14 and 16 show elevated *CD74* - associated interactions, identifying them as antigen-presentation niches. These results demonstrate EnsAgent helps explain functional heterogeneity within the tumor microenvironment.

### 3.3 Robustness to Batch Effects in Mouse Olfactory Bulb

The robustness of EnsAgent relies on the Critic-guided Rerun protocol embedded within the annotation module (Fig. 4a). During the initial phase, the Critic Agent detected a “Confidence Mismatch” where batch-induced noise compromised the Spatiality Expert. Instead of accepting unstable labels, the Orchestrator triggered a targeted re-evaluation using high-confidence inputs from Marker and Pathway experts. This second round successfully aligned the consensus identity and corrected errors before finalizing the output.

**Figure 4.**
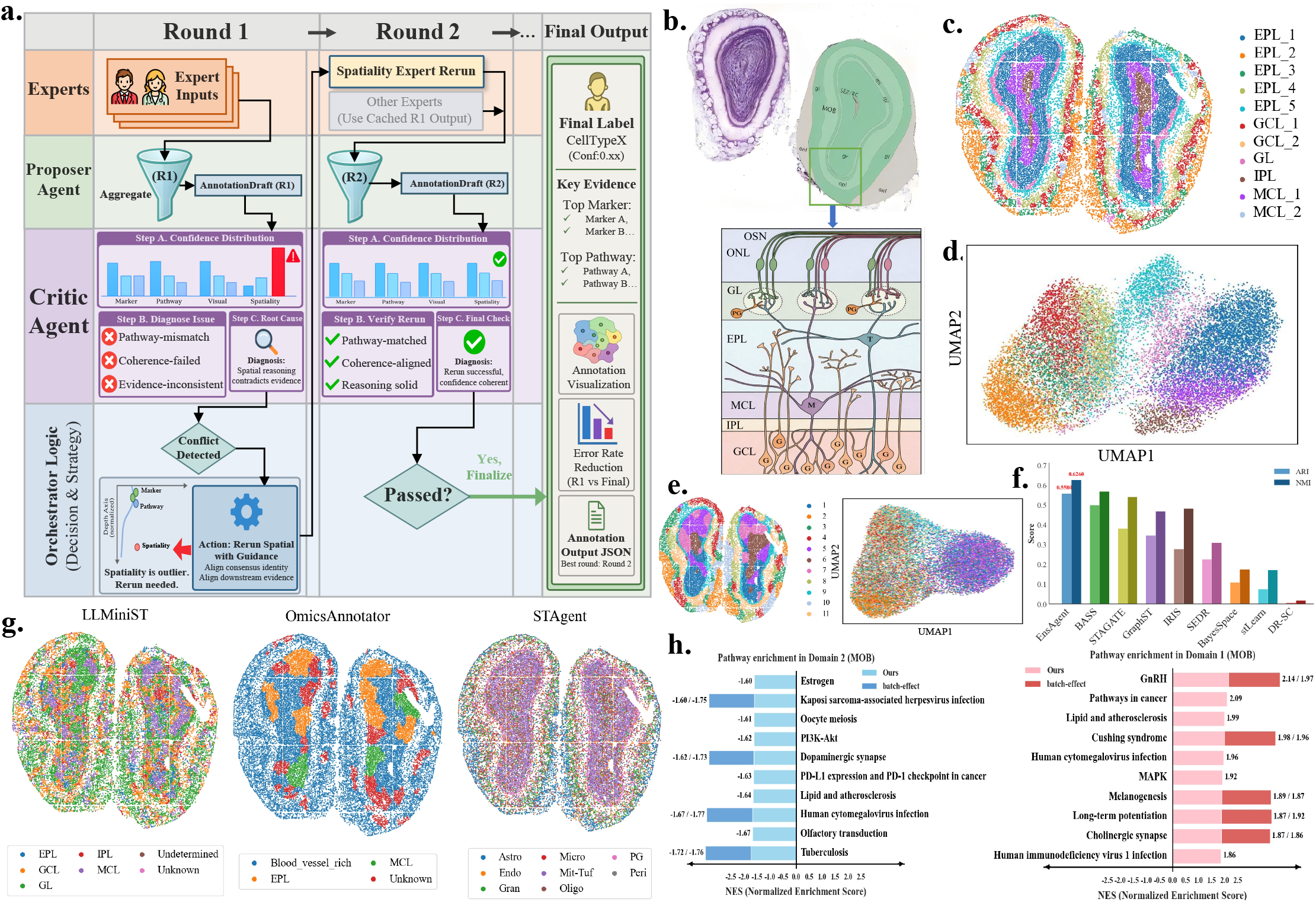
**(a)** Schematic of the Critic-guided Rerun protocol, where the agent actively corrects confidence mismatches. **(b)** Reference anatomical architecture (Allen Mouse Brain Atlas [Lein et al., 2007]), detailing layers from core to surface (SEL, GCL, IPL, MCL, EPL, GL, ONL). **(c)** EnsAgent domains recovering laminar organization under synthetic batch effects. **(d)** UMAP embedding of EnsAgent features showing batch neutralization. **(e)** Cluster results and UMAP visualization of single method baseline under batch effect noise conditions. **(f)** Quantitative robustness benchmarking (NMI/ARI). **(g)** Visual comparison vs. LLM-driven agents on MOB tissue. **(h)** Pathway enrichment analysis showing preservation of neuronal programs. **Alt Text:** (a) Rerun protocol; (b) MOB atlas with layers; (c) recovered domains; (d-e) UMAP scatter and spatial plots; (f) performance box plots; (g) spatial map comparison; (h) pathway charts.

This correction capability enables the recovery of a coherent laminar organization that aligns with the Allen Reference Atlas (Fig. 4b). Spatially, EnsAgent accurately delineated fine-grained sub-laminar compartments (Fig. 4c), capturing the subtle heterogeneity within the External Plexiform Layer (e.g., EPL_1–5_) and Granule Cell Layer (GCL_1–2_) while preserving the global superficial-to-deep trajectory from the Olfactory Nerve Layer (ONL) to the GCL.

To assess resilience to technical variation, we visualized the latent representations under synthetic batch effects. EnsAgent mitigated distortions through its multi-expert verification mechanism, yielding compact and topologically organized embeddings (Fig. 4d). In contrast, standard single-method pipelines produced markedly dispersed and intermixed latent UMAP representations (Fig. 4e), confirming vulnerability to batch-induced fragmentation (Supplementary Fig. S3). Quantitatively, EnsAgent achieved state-of-the-art agreement with reference labels, attaining an NMI of 0.6260 and ARI of 0.5580 while outperforming baseline clustering algorithms (Fig. 4f).

We further benchmarked anatomical fidelity against three representative LLM-driven agents on the raw MOB dataset (Fig. 4g). The baseline agents struggled to reconstruct the fine-grained laminar architecture. OmicsAnnotator yielded a coarse partition that failed to resolve the critical transition between the External Plexiform Layer(EPL) and Mitral Cell Layer(MCL). STAgent captured gross morphology but exhibited significant boundary leakage where the Granule Cell Layer erroneously extended into the Internal Plexiform Layer. Finally, LLMiniST produced highly fragmented annotations due to its lack of visual context. Its domains appeared as disjointed clusters rather than the continuous concentric rings characteristic of the olfactory bulb. Conversely, EnsAgent successfully reconstructed the intricate multi-layered organization validated by the atlas.

This structural fidelity grounds annotations in spatially resolved molecular evidence. Canonical markers identified by the Proposer Agent precisely trace the expected laminar organization (Supplementary Fig. S4), which translates into the recovery of functional interpretability (Fig. 4h). While technical noise attenuated key biological signals in single-method baselines, EnsAgent restored domain-specific programs. It recovered “MAPK signaling” in Domain 1 and the canonical “Olfactory transduction” alongside “PI3K-Akt” in Domain 2. Specifically, the specific enrichment of “MAPK signaling” in Domain 1 serves as a molecular hallmark that validates its annotation as the External Plexiform Layer (EPL), consistent with the established role of MAPK cascades in regulating synaptic plasticity at the dendrodendritic synapses characteristic of this region. These results confirm that Critic-guided refinement stabilizes spatial boundaries and rescues biologically meaningful functional programs from batch-induced distortions.

## 4 Discussion

We presented EnsAgent, a framework designed to decouple spatial domain annotation from the fragility of single-method partitioning. Unlike tool-augmented agents that passively orchestrate pipelines, or representation-based solvers that rely on fixed upstream inputs, EnsAgent introduces an active arbitration layer via its Scoring and Critic Agents. This shift from one-shot inference to a Consultation–Review workflow allows the system to filter technical noise before it propagates into semantic labels. As demonstrated in the Mouse Olfactory Bulb [Chen et al., 2022], our ensemble strategy effectively neutralizes batch-induced topology shifts that confound individual baselines, ensuring downstream annotation is grounded in biological signal rather than technical artifacts. Furthermore, the successful identification of “forgotten” tumor microenvironment subgroups in breast cancer tissue [Wu et al., 2021] highlights the critical value of our Proposer–Critic loop: the system does not merely assign a label but actively seeks evidence to justify it, effectively automating the “expert intuition” required to resolve conflict between data-driven clusters and biological expectations.

Despite these strengths, the parallel tool execution and iterative rerun cycles of EnsAgent entail higher computational overhead than single-paradigm baselines. Additionally, the current framework processes tissue sections individually and cannot yet leverage 3D spatial continuity from serial sections. To address latency, we propose using EnsAgent as a high-fidelity data generator to distill its ensemble reasoning into a specialized bio-spatial small language model (SLM) [Hsieh et al., 2023], yielding a lightweight inference engine. Concurrently, we aim to extend the architecture to support multi-slice alignment where agents query adjacent tissue layers for robust volumetric mapping.

## Author contributions

D.Z. and M.Z. conceived the method, developed the software, and wrote the original draft. N.L. and L.L. performed data curation, validation, and visualization. Q.D. and X.K. provided resources and supervision, and acquired funding. All authors contributed to formal analysis and reviewed the final manuscript.

## Supplementary data

Supplementary data are available at *Bioinformatics* online.

## Competing interests

No competing interest is declared.

## Notes

https://github.com/keviccz/ensAgent

### Competing Interest Statement

The authors have declared no competing interest.

