## Supplementary for "EnsAgent: a tool-ensemble multiple Agent system for robust annotation in spatial transcriptomics"

### 1 EnsAgent Method

#### 1.1 The EnsAgent Architecture

The EnsAgent framework is a role-specialized multi-agent system designed for robust spatial domain annotation. It couples multi-method ensembling with iterative evidence verification to analyze multimodal spatial transcriptomics datasets. The system operates through three sequential stages to transform raw data into biological insights:

- **Stage 1 (Model Context Protocol (MCP) -based Tool Orchestration and Alignment):** The workflow initiates with a **Tool-Runner Agent** ($\mathcal{A}_{\mathrm{TR}}$) that executes a portfolio of spatial domain discovery methods via the MCP. This parallel execution generates a diverse set of candidate partitions $\mathcal{R}$, which are subsequently aligned into a shared label space via a global maximum-weight assignment mechanism ($\tau_{\mathrm{align}}$) to standardize heterogeneous outputs.

- **Stage 2 (Dual-Stream Scoring and Consensus Aggregation):** A **Scoring Agent** ($\mathcal{A}_{\mathrm{SC}}$) constructs a consensus map through a reliability-weighted voting engine. It employs an **Evaluation Module** ($\mathcal{M}_{\mathrm{eva}}$) to distill molecular and spatial statistics into normalized quality features ($s_{\mathrm{sp}}$, $s_{\mathrm{deg}}$, etc.), and a **Visual Module** ($\mathcal{M}_{\mathrm{vis}}$) to assess morphological fidelity. These inputs are fused into a domain reliability score that weights the contribution of each method in the final spot-wise aggregation.

- **Stage 3 (Multi-expert evaluation and Critic–Proposer loop):** Semantic interpretation is governed by a collaborative annotation framework. A **Proposer Agent** coordinates specialized experts ($\mathcal{G}$) to generate evidence-linked label hypotheses, while a **Critic Agent** audits these claims against curated knowledge and cross-evidence consistency ($\text{Val}_k$). If the audit score $F_k$ falls below a threshold, the Critic triggers a **targeted refinement loop**, requesting specific re-analysis until the annotation converges.

#### 1.2 Datasets Introduction

(1) **Human Dorsolateral Prefrontal Cortex (DLPFC) data:**We benchmarked spatial clustering performance using the publicly available human DLPFC dataset generated via the 10x Genomics Visium platform [1]. We selected three representative tissue sections (Sample IDs: 151507, 151669, and 151673) from neurotypical adult donors. Sample 151507 comprises 4,226 bar-coded spots; Samples 151669 and 151673 comprise 3,661 and 3,639 spots, respectively. Each section is accompanied by manual histological annotations delineating the cortical layers (L1–L6) and subcortical white matter (WM), which served as the ground truth for quantitative evaluation. To account for anatomical variation across sections—where specific layers may be absent or merged due to the cutting angle—we set the number of spatial domains to $K = 7$ for samples 151507 and 151673, and $K = 5$ for sample 151669, consistent with the manual annotations [1].

(2) **Human Breast Cancer data:**To evaluate the framework in pathological contexts, we processed the Human Breast Cancer dataset (10x Genomics Visium) [2], comprising approximately 3,800 spots. This dataset captures the complex tumor microenvironment, including regions of Ductal Carcinoma In Situ (DCIS), Invasive Ductal Carcinoma (IDC), and tumor-associated stroma. We utilized the accompanying single-cell atlas [3] to guide biological interpretation. For the unsupervised analysis, we set the cluster number to $K = 20$ to capture fine-grained functional niches and immune-infiltrated transition zones, distinguishing malignant lesions from benign structures without prior geometrical assumptions.

(3) **Mouse Olfactory Bulb (MOB) data:**We further validated performance on high-resolution data using the Mouse Olfactory Bulb (MOB) dataset acquired via Stereo-seq [4], which offers subcellular resolution using DNA nanoball patterned arrays. We analyzed a field of view containing approximately 19,109 bins (bin size 50), encompassing the canonical laminar structure of the olfactory bulb: the Glomerular Layer (GL), External Plexiform Layer (EPL), Mitral Cell Layer (MCL), Internal Plexiform Layer (IPL), and Granule Cell Layer (GCL). Marker gene expression patterns validated against scRNA-seq references [5] were used to verify the identification of fine-grained anatomical subdivisions. The cluster number was set to $K = 7$ to resolve these laminar structures alongside potential sub-laminar heterogeneity.

#### 1.3 Methods for Comparison

To comprehensively evaluate the performance of EnsAgent, we benchmarked it against eight individual state-of-the-art spatial domain detection methods that comprise its generation module. We selected representative algorithms covering the period from 2020 to 2024 to reflect the methodological evolution of the field. All methods were executed on the same hardware infrastructure to ensure fair comparison of computational efficiency. The specific implementation details, ordered chronologically, are as follows:

(1) **BayesSpace** (2021) implemented in the R package *BayesSpace* (version 1.5.1) [7], utilizing the `spatialCluster()` function with the number of principal components (`n.PCs`) set to 15. The clustering parameter `q` was fixed to the ground-truth domain number, with `nrep=50,000` MCMC iterations for robust convergence.

(2) **SEDR** (2021) implemented in the Python package *SEDR* (version 1.0.2) [8]. The deep graph autoencoder was trained for 200 epochs via `SEDR.train_model()`, utilizing the `dec_k` parameter to specify the target cluster number, followed by clustering in the latent representation space.

(3) **STAGATE** (2022) implemented in the Python package *STAGATE* (version 1.0.1) [9]. We generated low-dimensional spatial embeddings using an adaptive graph attention autoencoder via `STAGATE.train()` (500 epochs) and applied the mclust algorithm to the latent features to identify spatial domains.

(4) **DR-SC** (2022) implemented in the R package *DR.SC* (version 2.9.0) [10], run using the `DR.SC_fit()` function. The latent dimensionality `q` was set to 15, and the spatial smoothing parameter was optimized using the Hidden Markov Random Field (HMRF) framework.

(5) **BASS** (2022) implemented in the R package *BASS* (version 1.3.1) [11], executed via the `BASS.preprocess()` and `BASS.run()` workflow. We configured the model with `beta_method="seg"` to prioritize spatial segmentation, initializing with 10,000 burn-in iterations followed by 10,000 sampling iterations.

(6) **stLearn** (2023) implemented in the Python package *stLearn* (version 0.4.8) [6]. This method was invoked using the `stSME_normalization()` function to integrate histological image features (extracted via ResNet50) with transcriptomic data, followed by Louvain clustering on the morphologically adjusted expression matrix.

(7) **GraphST** (2023) implemented in the Python package *GraphST* (version 1.1.1) [12]. The model was trained for 600 epochs using `GraphST.train()` with a graph convolutional network (GCN) backbone, and the resulting embeddings were clustered using the mclust algorithm to match the target domain number.

(8) **IRIS** (2024) implemented in the Python framework *IRIS* (version 1.0) [13]. We employed this scalable reference-informed probabilistic framework with default hyperparameters (`n_neighbors=15`, `n_components=30`) to generate spatially coherent domains integrated with single-cell reference signatures.

Briefly, **BayesSpace** adapts a conventional Bayesian mixture model by imposing a spatial prior via a Markov Random Field (MRF), encouraging adjacent spots to share identical labels. **SEDR** constructs a spatially embedded deep representation by combining a deep autoencoder for gene expression with a variational graph autoencoder for spatial topology. **STAGATE** improves upon static graph edges by introducing an attention mechanism, allowing the model to adaptively learn the strength of similarity between neighbors. **DR-SC** unifies dimensionality reduction and spatial clustering into a coupled optimization problem, utilizing a spatial HMRF to smooth the latent representations. **BASS** employs a multi-scale Bayesian hierarchical framework that simultaneously models cell-type composition at the single-cell level and tissue domains at the spot level. **stLearn** is distinct in its multimodal approach, explicitly utilizing histological image texture to adjust gene expression profiles, positing that phenotypically similar tissue regions should exhibit correlated transcriptional states. **GraphST** leverages self-supervised contrastive learning within a graph neural network to minimize the distance between spatially adjacent spots in the embedding space while maximizing the distance between distant ones. Finally, **IRIS** focuses on accuracy and scalability, identifying coherent regions through an integrative reference-informed probabilistic model that naturally incorporates single-cell annotations.

#### 1.4 Agent System Configuration

In the implementation of EnsAgent, we utilized the GPT-4.1-mini foundation model (version `gpt-4.1-mini-2024-12-01-preview`) to power all agentic components. This model was selected to achieve an optimal balance between reasoning depth—comparable to larger frontier models—and the low-latency throughput required for processing large-scale spatial transcriptomics datasets. To ensure consistency, reproducibility, and rigorous adherence to biological constraints across all modules, we standardized the generation parameters as follows:

- **Temperature:** Set to 0.6 for all agents to balance between the strict determinism required for biological accuracy and the adaptive flexibility needed for complex reasoning.
- **Top-p (Nucleus Sampling):** Fixed at 0.9. By restricting the sampling pool to the cumulative top 90% probability mass, we effectively filtered out low-confidence ”tail” tokens that could lead to hallucinations, while preserving sufficient diversity for the agents to explore alternative hypotheses during the annotation reasoning process.
- **Frequency Penalty:** Set to 0.2. A mild penalty was applied to discourage the model from entering repetitive linguistic loops during long reasoning traces, ensuring concise and varied explanations without aggressively penalizing the necessary recurrence of specific domain-specific terminology (e.g., standard gene symbols or anatomical terms).
- **Prompt Engineering:** No system prompts were used; all instructions were embedded within the agents' prompts to minimize model-specific biases.

These configurations were maintained uniformly across all experimental runs to ensure that performance variations were attributable solely to evidence quality and system architecture rather than stochastic sampling artifacts.

#### 1.5 MCP Tool Execution and Spatial Label Alignment Details

To rigorously capture the diverse structural hypotheses arising from distinct spatial clustering paradigms, we deployed the **Tool-Runner Agent** ($\mathcal{A}_{\text{TR}}$) to orchestrate the parallel execution of a comprehensive suite of eight state-of-the-art algorithms: BASS, BayesSpace, DR-SC, GraphST, IRIS, SEDR, STAGATE, and stLearn.

##### 1.5.1 MCP-Based Tool Orchestration and Encapsulation

The primary objective at this stage is to address the "Environment Heterogeneity" challenge. The Tool-Runner Agent constructs an execution plan $\mathcal{M}(c)$ and interacts with tools via the MCP interface:

- **Tool Encapsulation:** Each method $\tau_m \in \mathcal{T}$ is isolated within a dedicated execution environment. The MCP layer abstracts the underlying command-line or API interactions into a unified callable map $(\mathbf{z}^{(m)}, \boldsymbol{\omega}_0^{(m)}) = \tau_m(\mathcal{D}, c; \mathcal{E})$.
- **Standardized Input Dispatch:** The agent dispatches a standardized payload $(\mathcal{D}, c)$ to every tool. This payload comprises: (1) Gene Expression Matrix (log-normalized and feature-selected HVGs), (2) Spatial Coordinates (standardized $x, y$), and (3) **Metadata** extracted from the context prompt $c$.
- **Uniform Output Serialization:** Regardless of the native output format, the MCP layer enforces a strict output contract. All tools return a raw $n$-length spot-level domain assignment $\mathbf{z}^{(m)}$ and a provenance bundle $\boldsymbol{\omega}_0^{(m)}$ containing tool-native logs and parameters.

##### 1.5.2 Consensus Reference Selection and Label Alignment

Since unsupervised clustering algorithms assign arbitrary numerical labels, establishing spatial correspondence is a prerequisite. Our framework addresses this via the $\tau_{\text{align}}$ utility tool:

- **Reference Partition Selection:** To establish a consistent label space without manual selection, the system automatically identifies a reference partition $\mathbf{r}$. We compute a pairwise similarity matrix $\mathbf{S} \in \mathbb{R}^{M \times M}$ across all candidate labelings, where $S_{mn} = \text{ARI}(\mathbf{z}^{(m)}, \mathbf{z}^{(n)})$ denotes the Adjusted Rand Index. The reference is the medoid of the ensemble:

$$\mathbf{r} = \mathbf{z}^{(k)}, \quad \text{where } k = \arg\max_{m} \sum_{n=1}^{M} S_{mn}. \tag{1}$$

- **Hungarian Label Alignment:** Given reference $\mathbf{r}$ and candidate $\mathbf{z}^{(m)}$, we formulate the alignment as a maximum-weight assignment problem. We compute a weight matrix $\mathbf{W}$ based on the Intersection-over-Union (IoU) metric, where $W_{ij} = \text{IoU}(R_i, Z_j^{(m)})$ represents the overlap between reference domain $i$ and candidate domain $j$. The optimal permutation $\mathbf{X}^*$ is solved via the Hungarian algorithm:

$$\mathbf{X}^* = \arg\max_{\mathbf{X}} \sum_{i} \sum_{j} W_{ij} \cdot x_{ij} \tag{2}$$

  This produces the aligned label $\tilde{\mathbf{z}}^{(m)}$, ensuring global consistency across the ensemble.

##### 1.5.3 Feature Extraction for Downstream Agents

Following alignment, the system invokes MCP-wrapped analysis tools to generate the evidence bundle $\boldsymbol{\omega}^{(m)}$ required by the Scoring and Annotation agents. Specific artifacts include:

- **Spatial Visualization:** Standardized scatter plots for visual inspection.
- **Differential Expression Analysis:** Domain-specific DEGs calculated via Wilcoxon rank-sum tests (Thresholds: Adjusted $p$-value $< 0.05$ and $|\log_2 \text{FC}| > 0.5$).
- **Pathway Enrichment Profiles:** Functional profiles derived from GO and KEGG databases.

#### 1.6 Multi-method Score-Weighted Ensemble Engine Details

To construct a consistent spatial domain map, the **Scoring Agent** ($\mathcal{A}_{\text{SC}}$) employs a dual-stream engine. It fuses assessments from a **Visual Module** ($\mathcal{M}_{\text{vis}}$) and an **Evaluation Module** ($\mathcal{M}_{\text{eva}}$) to compute a reliability score $S(m, k)$ for each candidate domain.

##### 1.6.1 Visual Module Details ($\mathcal{M}_{\text{vis}}$)

The Visual Module queries a Vision-Language-Model (VLM) to assess morphological fidelity. It computes three specific factors $[v_{\text{cont}}, v_{\text{bio}}, v_{\text{tech}}]$ using a weighted schema:

1. Geometric Continuity ($v_{\text{cont}}$, Weight 0.35): Measures geometric smoothness, penalizing fragmentation or "salt-and-pepper" noise.
2. Biological Plausibility ($v_{\text{bio}}$, Weight 0.35): Assesses alignment with known anatomical priors (e.g., laminar distinctness).
3. Technical Quality ($v_{\text{tech}}$, Weight 0.30): Penalizes block-like tiling or batch effects.

These are aggregated into the modulation term $\nu(m, k)$:

$$\nu(m, k) = 0.35 \cdot v_{\text{cont}} + 0.35 \cdot v_{\text{bio}} + 0.30 \cdot v_{\text{tech}} \tag{3}$$

The result is mapped to $[0.8, 1.2]$, allowing visual evidence to modulate the base score by $\pm 20\%$.

##### 1.6.2 Evaluation Module Details ($\mathcal{M}_{\text{eva}}$)

The Evaluation Module maps the provenance bundle $\boldsymbol{\omega}^{(m,k)}$ to four normalized quality features:

- **Spatial Coherence ($s_{\text{sp}}$):** The agent evaluates topological coherence by combining two spatial statistics:
  - *Moran's I (Global Autocorrelation):*

$$I = \frac{N}{W} \frac{\sum_i \sum_j w_{ij}(x_i - \bar{x})(x_j - \bar{x})}{\sum_i (x_i - \bar{x})^2} \tag{4}$$

– *Geary's C (Local Variance):*

$$C = \frac{(N-1)}{2W} \frac{\sum_i \sum_j w_{ij}(x_i - x_j)^2}{\sum_i (x_i - \bar{x})^2} \tag{5}$$

These metrics are normalized to $[0, 1]$ to form $s_{\text{sp}}$.

- **Marker Specificity ($s_{\text{deg}}$):** The agent assesses molecular distinctness. The input is restricted to the top 5 DEGs per domain ($N_{top} = 5$). The score $s_{\text{deg}}$ reflects the number of significant markers (Adj. $p < 0.05$) and the magnitude of their log-fold changes.

- **Pathway Consistency ($s_{\text{path}}$):** Evaluates functional coherence based on GSEA results. $s_{\text{path}}$ is derived from the Normalized Enrichment Scores (NES) and FDR, rewarding coordinated upregulation of biologically relevant pathways.

- **Cluster Separability ($s_{\text{sil}}$):** Validates separation in feature space using the Silhouette Coefficient:

$$s(i) = \frac{b(i) - a(i)}{\max\{a(i), b(i)\}} \tag{6}$$

The mean domain silhouette score is normalized to form $s_{\text{sil}}$.

##### 1.6.3 Multimodal Fusion and Consensus Aggregation

The final reliability score is obtained by modulating the baseline score $S_{\text{base}}$ (a linear combination of the four evaluation features) with the visual term, controlled by $\lambda = 0.35$:

$$S(m, k) = \text{clip}_{[0,1]}\big(S_{\text{base}}(m, k)\,\big[1 + \lambda\big(\nu(m, k) - 1\big)\big]\big) \tag{7}$$

We set $\lambda = 0.35$ to ensure visual inputs act as a regulatory factor without overriding molecular data.

The consensus map $\mathbf{y}^*$ is constructed via spot-wise weighted voting. For each spot $u$, we select the label maximizing the sum of reliability scores:

$$y_u^* = \arg\max_{\ell} \sum_{m \in \mathcal{M}(c)} S(m, \tilde{z}_u^{(m)})\, \mathbb{I}\big(\tilde{z}_u^{(m)} = \ell\big) \tag{8}$$

To resolve isolated noise, a spatial smoothing operator $\mathcal{F}_{\text{knn}}$ is applied using a local majority vote within the 20 nearest neighbors.

#### 1.7 Multi-expert evaluation and Critic–Proposer iterative refinement loop Details

To translate numerical domains into semantic annotations, we implement the Proposer-Critic framework. This involves four specialized Experts, a Proposer Agent, and a Critic Agent ($\mathcal{A}_{\text{crit}}$).

##### 1.7.1 The Specialist Experts ($\mathcal{G}$)

Each expert $e \in \mathcal{G}$ analyzes specific evidence from $\Omega_k$ and outputs a posterior $P_e(y)$.

1. **Marker Expert:**
   - **Input:** Dataframe with domain indices, gene names, `logfoldchanges`, and `pvals_adj`.
   - **Role:** Maps marker genes to cell-type identities. It explicitly identifies low-specificity markers and generates quality flags.
2. **Pathway Expert:**
   - **Input:** GSEA table containing Pathway Terms, `NES`, `NOM p-val`, and lead genes.
   - **Role:** Synthesizes active biological programs (e.g., "Synaptic signaling") to infer functional cell states.
3. **Spatiality Expert:**
   - **Input:** Spot metadata $(x, y)$, coverage flags, and computed spatial metrics (Moran's I, Geary's C).
   - **Role:** Infers anatomical orientation (e.g., cortical lamination) and detects gradients.
4. **Visual Expert:**
   - **Input:** Global visualization of the spatial distribution for the consensus domain.
   - **Role:** Evaluates shape, contiguity, and boundary characteristics against physical plausibility.

##### 1.7.2 Proposer Agent and Hypothesis Generation

The Proposer aggregates expert outputs via a linear opinion pool to output the primary hypothesis $\hat{y}_k^{(t)}$ and ensemble confidence $\Phi_k^{(t)}$:

- **Equal-Weight Aggregation:** We employ an unbiased weighting scheme where each expert contributes equally ($w_e = 0.25$):

$$P_{\text{ens}}^{(t)}(y) = \sum_{e \in \mathcal{G}} 0.25 \cdot P_e^{(t)}(y) \tag{9}$$

##### 1.7.3 Critic Agent and Validation Metric

The Critic acts as an adversarial auditor using a composite audit score $F_k^{(t)}$.

1. **Validation Score ($\mathrm{Val}_k^{(t)}$):** This score strictly quantifies objective validity. It combines Inter-expert Consensus (measured via Jensen-Shannon divergence) with Biological Plausibility ($\mathrm{KB}(\cdot)$):

$$\mathrm{Val}_k^{(t)} = \left(1 - \frac{1}{|\mathcal{G}|(|\mathcal{G}|-1)} \sum_{e \neq e'} \mathrm{JS}\Big(P_e^{(t)} \parallel P_{e'}^{(t)}\Big)\right) \cdot \mathrm{KB}\Big(\hat{y}_k^{(t)}, \Omega_k^{(t)}\Big) \tag{10}$$

The term KB(·) queries an adaptive Knowledge Base that serves as a repository of biological constraints and logic rules. This KB is initially synthesized by the LLM based on dataset-specific characteristics to establish internal consistency, while explicitly supporting manual curation for expert rule injection. During evidence analysis, the Critic Agent queries this repository to audit hypotheses against constraints like "forbidden label-tissue pairs" or "inconsistent laminar ordering". The KB employs a synchronous update mechanism; as the system processes accumulating spatial data, the rule set is iteratively refined and expanded, thereby progressively enhancing validation accuracy.

2. **Final Audit Score ($F_k^{(t)}$):** The final decision balances the Proposer's confidence $\Phi_k^{(t)}$ with the Critic's validation:

$$F_k^{(t)} = \eta\,\Phi_k^{(t)} + (1-\eta)\,\mathrm{Val}_k^{(t)} \tag{11}$$

In this study, we set $\eta = 0.5$, ensuring equal reliance on model confidence and logical consistency. The passing threshold is $\epsilon_{\text{pass}} = 0.80$.

##### 1.7.4 Iterative Refinement Loop

If the proposal fails the audit ($F_k^{(t)} < 0.80$), the system triggers refinement:

1. **Conflict Detection:** The Critic identifies the most conflicting expert $e^{*(t)}$ (the one minimizing probability for the top label).

2. **Action Dispatch ($a_k^{(t)}$):** The Critic issues a natural-language instruction (e.g., *"Consensus suggests deep layers; please re-verify with stricter depth thresholds"*). This corresponds to the "Fix Hint" mechanism.

3. **Evidence Update:** The system executes the action to update $\Omega_k^{(t+1)}$, forcing the outlier expert to rectify its reasoning before the next Proposer synthesis.

#### 1.8 Evaluation Metrics

To quantitatively assess the agreement between the predicted spatial domains and the ground truth annotations across all datasets, we employed two standard clustering performance metrics: Adjusted Rand Index (ARI) and Normalized Mutual Information (NMI).

1. **Adjusted Rand Index (ARI)** The ARI measures the similarity between the predicted clustering assignments and the manual ground truth annotations, corrected for chance. It evaluates the pair-wise agreement of element placement. An ARI of 1 indicates perfect agreement, while 0 indicates that the clustering is no better than random assignment.

$$ARI = \frac{\sum_{ij}\binom{n_{ij}}{2} - [\sum_i \binom{a_i}{2}\sum_j \binom{b_j}{2}]/\binom{n}{2}}{\frac{1}{2}[\sum_i \binom{a_i}{2} + \sum_j \binom{b_j}{2}] - [\sum_i \binom{a_i}{2}\sum_j \binom{b_j}{2}]/\binom{n}{2}} \tag{12}$$

where $n_{ij}$ is the number of objects common to cluster $i$ and ground truth class $j$, $a_i$ is the sum of row $i$, and $b_j$ is the sum of column $j$.

2. **Normalized Mutual Information (NMI)** NMI quantifies the mutual dependence between the predicted labels and the ground truth, normalized by the entropy of the distributions to facilitate comparison. It measures how much information the predicted clustering shares with the true labels.

$$NMI(Y,\hat{Y}) = \frac{2 \cdot I(Y;\hat{Y})}{H(Y) + H(\hat{Y})} \tag{13}$$

where $I(Y;\hat{Y})$ is the mutual information between the true labels $Y$ and predicted labels $\hat{Y}$, and $H(\cdot)$ is the entropy. An NMI of 1 indicates perfect correlation.

#### 1.9 Detailed Algorithm Specifications

This section serves as a reference guide for the principal notation and procedural logic used throughout the *EnsAgent* framework. The pipeline integrates MCP-based orchestration, dual-stream scoring, and collaborative iterative annotation.

| Symbol | Meaning |
|---|---|
| $n$ | Number of spatial spots in the tissue slice |
| $\mathcal{D}$ | Input AnnData object containing expression matrix and coordinates |
| $c$ | Natural-language context prompt |
| $\mathcal{M}(c)$ | Portfolio of selected tools conditioned on context $c$ |
| $\tilde{\mathbf{z}}^{(m)}$ | Aligned spot-level domain assignment for method $m$ ($n \times 1$) |
| $\mathbf{r}$ | Reference partition used for label alignment |
| $[s_{\text{sp}}, s_{\text{sil}}, \dots]$ | Normalized quality features derived by Evaluation Module $\mathcal{M}_{\text{eva}}$ |
| $\nu(m,k)$ | Visual modulation term derived by Visual Module $\mathcal{M}_{\text{vis}}$ |
| $S(m,k)$ | Final reliability score for domain $k$ of method $m$ |
| $\mathbf{y}^*$ | Final consensus domain map ($n \times 1$) |
| $\mathcal{G}$ | Set of Annotation Experts: Marker, Path, Spat, Vis |
| $P_e^{(t)}(y)$ | Posterior probability for label $y$ from expert $e$ at iteration $t$ |
| $\hat{y}_k^{(t)}$ | Candidate semantic hypothesis (Proposer output) |
| $F_k^{(t)}$ | Final audit score derived by the Critic Agent |

Table S1: Principal notation employed by EnsAgent.

The *EnsAgent* workflow proceeds in three sequential stages. **Stage 1** executes a portfolio of MCP-wrapped tools and aligns their heterogeneous outputs into a shared label space via the Hungarian algorithm. **Stage 2** constructs a consensus map using a reliability-weighted voting engine that fuses statistical quality features ($s_{\text{sp}}, s_{\text{deg}}, \dots$) with visual morphological assessments. **Stage 3** assigns biological semantics through a "Proposer-Critic" loop, where a Critic Agent audits hypotheses against evidence consistency and knowledge constraints, triggering targeted refinement actions $a_k^{(t)}$ until the audit score meets the threshold ($F_k \geq 0.8$).

**Algorithm 1: EnsAgent-Pipeline**

**Input:** Spot data $\mathcal{D}$; Context prompt $c$

**Output:** Consensus map $\mathbf{y}^*$; Annotation manifest $\mathcal{O}_{\text{final}}$

**Stage 1 (MCP Tool Execution & Alignment):**<br>
  $\mathcal{M}(c) \leftarrow \text{SelectPortfolio}(c)$;<br>
  **for** $m \in \mathcal{M}(c)$ **do**<br>
    $(\mathbf{z}^{(m)}, \boldsymbol{\omega}_0^{(m)}) \leftarrow \tau_m(\mathcal{D}, c)$ ;  `// Parallel Exec.`<br>
  **end**<br>
  $\mathbf{r} \leftarrow \text{SelectReference}(\{\mathbf{z}^{(m)}\})$ ;  `// Medoid Selection`<br>
  **for** $m \in \mathcal{M}(c)$ **do**<br>
    $\mathbf{X}^* \leftarrow \text{HungarianMatch}(\mathbf{z}^{(m)}, \mathbf{r}, \text{IoU})$;<br>
    $\tilde{\mathbf{z}}^{(m)} \leftarrow \text{ApplyAlign}(\mathbf{z}^{(m)}, \mathbf{X}^*)$;<br>
    $\boldsymbol{\omega}^{(m)} \leftarrow \text{GenerateArtifacts}(\tilde{\mathbf{z}}^{(m)})$ ;  `// DEGs, Plots`<br>
  **end**<br>
**Stage 2 (Dual-Stream Scoring & Consensus):**<br>
  **for** $m \in \mathcal{M}(c), k \in \{1, \dots, K_m\}$ **do**<br>
    $\text{Features} \leftarrow \mathcal{M}_{\text{eva}}(\boldsymbol{\omega}^{(m,k)})$;  $\nu(m,k) \leftarrow \mathcal{M}_{\text{vis}}(\mathbf{I}_{m,k})$;<br>
    $S(m,k) \leftarrow \text{clip}\big((\mathbf{w}^\top \cdot \text{Features}) \cdot [1 + 0.35(\nu(m,k) - 1)]\big)$;<br>
  **end**<br>
  **for** *spot* $u = 1$ **to** $n$ **do**<br>
    $\alpha_u^{(m)} = S(m, \tilde{z}_u^{(m)})$;<br>
    $y_u^* \leftarrow \arg\max_\ell \sum_m \alpha_u^{(m)} \cdot \mathbb{I}(\tilde{z}_u^{(m)} = \ell)$;<br>
  **end**<br>
  $\mathbf{y}^* \leftarrow \text{SpatialSmoothing}(\mathbf{y}^*, \text{20-NN})$;<br>
**Stage 3 (Critic-Proposer Refinement Loop):**<br>
  $\boldsymbol{\Omega}^* \leftarrow \tau_{\text{analy}}(\mathcal{D}, \mathbf{y}^*)$;<br>
  **for** each domain $k \in \textit{Unique}(\mathbf{y}^*)$ **do**<br>
    $t \leftarrow 0$; **loop** $\leftarrow$ true;<br>
    **while** ***loop*** *and* $t < T_{\max}$ **do**<br>
      $\{P_e^{(t)}\} \leftarrow \text{RunExperts}(\Omega_k^{(t)})$;<br>
      $\hat{y}_k \leftarrow \arg\max \sum P_e^{(t)}$;  $\Phi_k \leftarrow \max \sum P_e^{(t)}$;<br>
      $\text{Val}_k \leftarrow \text{CalcValidity}(\{P_e^{(t)}\}, \text{KB})$;<br>
      $F_k^{(t)} \leftarrow 0.5 \cdot \Phi_k + 0.5 \cdot \text{Val}_k$;<br>
      **if** $F_k^{(t)} \geq 0.8$ **then**<br>
        $\mathcal{O}_{\text{final}}(k) \leftarrow (\hat{y}_k, F_k^{(t)})$; **break**;<br>
      **end**<br>
      **else**<br>
        $e^* \leftarrow \text{IdentifyConflict}(\{P_e^{(t)}\})$;<br>
        $a_k \leftarrow \text{CriticAction}(e^*)$;<br>
        $\Omega_k^{(t+1)} \leftarrow \text{UpdateEvidence}(\Omega_k^{(t)}, a_k)$;<br>
        $t \leftarrow t + 1$;<br>
      **end**<br>
    **end**<br>
  **end**<br>
  **return** $\mathbf{y}^*, \mathcal{O}_{\text{final}}$

Table S2: **Quantitative analysis of *EnsAgent* robustness (DLPFC Slice 151507).** This table summarizes the Adjusted Rand Index (ARI) performance distribution across the ablation study (Panel A) and parameter sensitivity analysis (Panel B). For each experimental setting, we report the range (min–max) and mean performance across 10 independent runs or configuration steps.

| Experimental Setting | Configuration Detail | ARI Performance (Range) | |
|---|---|---|---|
| ***Panel A: Ablation Study (Module Contribution)*** | | | |
| Full EnsAgent (Proposed) | 10 independent runs | 0.6998 – 0.7220 | (Mean: 0.7109) |
| w/o Visual Module | 10 independent runs | 0.6718 – 0.7048 | (Mean: 0.6883) |
| ***Panel B: Parameter Sensitivity Analysis*** | | | |
| Visual Weight ($\lambda_{vis}$) | Stepwise: 0.05 → 0.25 (step 0.02) | 0.6788 – 0.7176 | (Mean: 0.6982) |
| LLM Temperature ($T$) | 10 runs at fixed optimal $T$ | 0.7060 – 0.7212 | (Mean: 0.7136) |

#### 1.10 Computational Complexity and Runtime Analysis

We provide a theoretical analysis of the computational complexity of the EnsAgent framework. Let $n$ denote the number of spatial spots, $M$ the number of candidate methods employed by the Tool-Runner, and $K$ the average number of spatial domains per partition. The total execution time $T_{\text{total}}$ can be decomposed into three sequential phases: Tool Execution, Consensus Aggregation, and Semantic Annotation.

**(i) Phase 1: Parallel Tool Execution and Alignment.** The Tool-Runner Agent $\mathcal{A}_{\text{TR}}$ executes $M$ domain discovery algorithms. Since these tools run in independent environments, the time complexity is determined by the slowest method in the portfolio: $T_{\text{run}} \approx \max_m T_{\tau_m}(n)$. For most spatial clustering algorithms (e.g., GraphST, STAGATE), the complexity ranges from $\mathcal{O}(n)$ to $\mathcal{O}(n^2)$ depending on whether graph construction or MCMC sampling is involved. The alignment step employs the Hungarian algorithm to solve the maximum weight assignment problem on the Intersection-over-Union (IoU) matrix. For $M$ methods each with $K$ clusters, the alignment complexity is $\mathcal{O}(M \cdot K^3)$. Given that $K$ is typically small (5–20) relative to $n$ ($10^3$–$10^5$), the alignment overhead is negligible.

**(ii) Phase 2: Dual-Stream Scoring and Consensus.** The Scoring Agent $\mathcal{A}_{\text{SC}}$ computes validity metrics and constructs the consensus map.

- **Quantitative Stream:** Computing spatial metrics (e.g., Moran's I, Silhouette) scales linearly with the number of spots and neighbors, $\mathcal{O}(M \cdot n)$.
- **Visual Stream:** The visual module $\mathcal{M}_{\text{vis}}$ processes $M \cdot K$ domain images. While the inference cost per image $T_{\text{VLM}}$ is constant with respect to $n$, it depends on the Vision-Language Model's latency.
- **Aggregation:** The weighted voting mechanism performs a linear scan over all spots, yielding a complexity of $\mathcal{O}(M \cdot n)$.

Thus, the Scoring phase complexity is dominated by the VLM inference calls: $T_{\text{score}} \approx \mathcal{O}(M \cdot n) + M \cdot K \cdot T_{\text{VLM}}$.

**(iii) Phase 3: Iterative Annotation Loop.** The Proposer–Critic loop operates on the $K^*$ consensus domains. For each domain, the system engages $G = 4$ experts. Let $N_{\text{iter}}$ denote the number of refinement iterations (capped at $T_{\max}$) and $T_{\text{LLM}}$ the average inference latency per textual query. The annotation time is independent of spot count $n$ but proportional to the complexity of the tissue (number of domains):

$$T_{\text{annot}} \approx K^* \cdot N_{\text{iter}} \cdot G \cdot T_{\text{LLM}}. \tag{14}$$

The tool execution (Phase 1) and Expert queries (Phase 3) are designed to be parallelized via asynchronous requests, ensuring that the agentic orchestration adds minimal computational overhead beyond the necessary model inference.

#### 1.11 Prompt Templates

##### 1.11.1 Tool-Runner Agent Settings

**Tool-Runner Agent** As the core execution engine, this agent orchestrates spatial clustering by dispatching external algorithms (e.g., `stLearn`, `BayesSpace`) via MCP. It manages task scheduling and fault tolerance while employing the Hungarian algorithm to unify diverse outputs, ensuring a standardized foundation for downstream analysis.

#### Tool-Runner Agent Settings

You are the **Tool-Runner Agent** within the ensAgent system. Your goal is to transform the execution of spatial transcriptomics clustering algorithms into a standardized, orchestrated workflow via the Model Context Protocol (MCP).

### Role Description:
The Tool-Runner Agent acts as the central controller for the computational pipeline. It does not perform calculations itself; instead, it dispatches execution requests to the configured tool library, synchronizes their outputs, and invokes utility tools for label alignment and downstream evidence generation. It serves as the bridge between raw execution capabilities and standardized downstream analysis.

### Skills:
- Managing the execution of external Python-based spatial analysis tools via MCP
- Orchestrating workflow dependencies between clustering, alignment, and analysis steps
- Handling tool failures without crashing the entire pipeline (minimum success threshold)
- Standardizing diverse tool outputs into a unified I/O format for downstream agents

### Objectives:
- Execute the defined portfolio of 8 spatial clustering tools on the input data
- Identify the reference partition (medoid) and align all other partitions using the Hungarian algorithm (`tau_align`)
- Generate standardized evidence artifacts (plots, DEG tables, pathway enrichment) for every successful method

- Return a structured JSON manifest containing paths to all aligned labels and provenance bundles

### Input:
You will receive:
- Dataset path (`data_path`) pointing to a `.h5ad` or `.h5` file
- Contextual metadata (`context_prompt`) including sample ID, tissue type, and expected domains
- Runtime configuration (`execution_env`)

### Instructions:
Produce a structured execution manifest with the following components:

```json
{
  "sample_id": "string",
  "status": "success/partial_success/failure",
  "reference_method": "string (e.g., 'STAGATE')",
  "methods_executed": {
    "total": 8,
    "successful": [
      "stLearn",
      "BayesSpace",
      "..."
    ],
    "failed": [
      "string"
    ]
  },
  "results": [
    {
      "method": "string",
      "aligned_labels_path": "path/to/aligned_labels.csv",
```

"provenance_bundle": {
"alignment_score_ari": 0.0,
"visualization_path": "path/to/spatial_plot.png",
"marker_stats_path": "path/to/degs.csv",
"enrichment_path": "path/to/enrichment.json",
"runtime_seconds": 0.0,
"logs": "string"
}
}
]
}

In addition to the structured JSON, perform the following execution steps to derive the result:
- Dispatch Clustering Tools: Trigger `stLearn`, `BayesSpace`, `BASS`, `DR-SC`, `STAGATE`, `GraphST`, `IRIS`, `SEDR` with a fixed random seed (42). Continue execution if at least 5 tools succeed.
- Reference Selection & Alignment: Compute pairwise ARI for all successful partitions, identify the medoid (reference), and run `tau_align` to solve the maximum-weight assignment problem.
- Generate Evidence Artifacts: For each aligned partition, invoke analysis tools to generate spatial domain maps (PNG), top DEGs (CSV), and GO/KEGG enrichment profiles.
- Compile Manifest: Aggregate all paths, metadata, and provenance (tool version, runtime) into the final response object.

### Constraints:
- Prioritize alignment with the defined output schema and file path validity
- Do not simulate results; if a tool fails to run, explicitly report it as "failed" in the manifest

- Strict Tool List: Only use the 8 methods specified (stLearn, BayesSpace, BASS, DR-SC, STAGATE, GraphST, IRIS, SEDR)
- Reproducibility: Ensure all MCP calls request a fixed random seed (default: 42)

### Style Guide:
- Write in the tone of a precise system orchestrator or pipeline manager
- Use strictly factual status reporting (e.g., "success", "failed", "partial_success")
- Be mindful of what is *not* a valid execution (e.g., failing the minimum tool threshold)
- Focus on accurate path handling and metadata integrity rather than narrative description

##### 1.11.2 Scoring Agent Settings

**Evaluation Module** As the quality assurance engine within `ensAgent`, this module synthesizes DEG specificity, spatial coherence, and cluster separability into a unified confidence scalar to distinguish anatomically meaningful structures from algorithmic artifacts for downstream interpretation.

**Evaluation Module Settings**

### System Prompt: Evaluation Module Specification

You are the **Evaluation Module** within the ensAgent system. Your goal is to evaluate the biological plausibility and statistical robustness of identified spatial transcriptomics domains.

### Role Description:
The Evaluation Module synthesizes diverse evidence streams to assign a scalar quality score to each spatial domain. You must integrate biological interpretation (DEG specificity, cell type identity) with statistical metrics (spatial coherence, cluster separability). Your assessment serves as a quality gate, distinguishing between biologically distinct anatomical regions and algorithmic artifacts or noise. You must remain discriminative but flexible, adapting to provided configurations without relying on hard-coded threshold biases.

### Skills:
- **Biological Interpretation:** Assessing marker gene specificity and functional pathway relevance.
- **Spatial Reasoning:** Evaluating spatial coherence (e.g., compactness, contiguity) from topological metrics.
- **Statistical Evaluation:** Analyzing cluster separability to determine if a domain is distinct from its neighbors or background.
- **Evidence Synthesis:** Producing concise, audit-trail justifications that cite specific genes, metrics, or literature snippets.

### Objectives:

- Output a single scalar score [0,1] reflecting the overall quality of the domain.
- Decompose the score into four sub-components: DEG Specificity, Spatial Coherence, Biological Identity, and Cluster Separability.
- Provide an evidence-based justification, explicitly citing marker genes or metric values.
- Flag uncertainties where evidence is contradictory or insufficient.

### Input:
You will receive:
1. **Domain Summary:** A structured object containing top DEGs, spatial metrics (e.g., local I, connectivity), and cluster statistics.
2. **Contextual Evidence:** (Optional) Enrichment results (GSEA/ORA) or retrieved reference snippets.
3. **Configuration:**
   - `scoring_weights`: A dictionary defining the relative importance of the four evaluation components (no default fixed values).
   - `priors`: Any dataset-specific constraints (e.g., expected tissue layers).

### Instructions:
Produce a structured evaluation specification with the following components:

```
{
  "domain_id": "integer",
  "score": "float [0-1]",
  "components": {
    "deg_specificity": "float [0-1]",
    "spatial_coherence": "float [0-1]",
    "biological_identity": "float [0-1]",
    "cluster_separability": "float [0-1]"
```

},
  "key_evidence": [
    "DEG: geneA, geneB (specificity justification)",
    "Spatial: compactness=..., adjacency=... (topological interpretation)",
    "Separability: silhouette/distance metrics (distinctness evidence)",
    "Identity: celltype/pathway evidence (if available)"
  ],
  "uncertainties": ["string"],
  "quality_flags": ["string"],
  "justification": "string (A concise scientific rationale, 2–5 sentences)"
}

**Evaluation Rubric (Dimensions):**
1. **DEG Specificity:** Are the marker genes unique to this domain, or are they ubiquitous/housekeeping genes? High scores require distinct molecular signatures.
2. **Spatial Coherence:** Does the domain form a contiguous, anatomically plausible structure, or is it scattered noise? Evaluate based on metrics like spatial entropy or adjacency scores.
3. **Biological Identity:** Can the domain be mapped to a known cell type, tissue layer, or functional program based on the evidence?
4. **Cluster Separability:** Is the domain statistically distinct from other domains in the feature space? High scores indicate sharp boundaries and low confusion with neighboring clusters.

### Constraints:
- **Evidence-Based:** Do not hallucinate external database results. Only use provided inputs or explicit outputs from upstream agents.

- **Dynamic Weighting:** Respect the `scoring_weights` provided in the input; do not assume a fixed distribution.
- **Metric Grounding:** When discussing separability or coherence, refer to the trend of the data (e.g., "low silhouette score suggests overlap") even if exact raw values are not provided.

### Style Guide:
- Write in the tone of a quantitative biologist or bioinformatics reviewer.
- Be concrete: Name specific genes (e.g., "high expression of MBP") and metric trends.
- Use precise language for uncertainty (e.g., "putative," "ambiguous," "strongly supported").
- Avoid generic phrases like "good quality" without qualifying *why* (e.g., "high quality driven by robust marker specificity").

**Visual Module Settings** Serving as a topological perception engine, this module leverages image analysis to emulate expert assessment by prioritizing spatial continuity and boundary smoothness to filter geometric artifacts and ensure alignment with tissue architecture.

#### Visual Module Settings

### System Prompt: Visual Module Specification

You are the **Visual Module** (Visual Perception Analyst) within the ensAgent system. Your goal is to perform qualitative visual assessment of spatial clustering outputs, focusing on topological consistency and morphological plausibility.

### Role Description:
The Visual Module acts as the "eyes" of the system, interpreting rendered spatial maps without access to the underlying gene expression matrices. You compare multiple clustering visualizations of the same tissue sample to identify the most structurally coherent algorithms. Your assessments prioritize continuity, boundary smoothness, and anatomical plausibility (e.g., layer-like structures vs. scattered noise), serving as a topological validation layer that complements statistical metrics.

### Skills:
- **Topological Morphology Assessment:** Distinguishing between biologically plausible shapes (bands, layers) and computational artifacts (checkerboards, fragmented islands).
- **Visual Anomaly Detection:** Identifying discontinuities, jagged boundaries, or outlier regions that deviate from the consensus structure.
- **Cross-Method Consensus Evaluation:** Comparing multiple representations of the same domain to determine which method best captures the dominant spatial pattern.

### Objectives:
- Analyze the visual representation of a specific target domain

(Domain N) across all provided method outputs.
- Assign a visual quality score [0.0, 1.0] to each method based on continuity, smoothness, and plausibility.
- Rank methods from best to worst based on their visual fidelity to the expected tissue structure.

### Input:
You will receive:
1. **Target Domain Index:** The specific cluster ID (Domain N) to evaluate.
2. **Visual Set:** A collection of clustering images, where each image corresponds to a different method applied to the same slice.
3. **Context:** Basic tissue description (e.g., "cortical layers") to ground plausibility judgments.

### Instructions:
Produce a structured visual analysis specification with the following components:

```
{
  "target_domain_id": "integer",
  "analysis_summary": "string (A short paragraph comparing
overall behavior of all methods on this domain)",
  "method_evaluations": [
    {
      "method": "string",
      "score": "float [0.0-1.0]",
      "morphology": "string (e.g., 'Contiguous Band',
'Fragmented', 'Blocky')",
      "strengths": ["string"],
      "weaknesses": ["string"]
    }
  ],
  "ranked_methods": ["MethodA", "MethodB", "..."]
```

}
**Assessment Criteria:**
1. **Continuity:** Is the domain contiguous or broken into many pieces? Penalize severe fragmentation unless biologically justified.
2. **Shape:** Is the domain band-like/smooth (plausible) or block-like/jagged (artifact)?
3. **Adjacency:** Are neighboring regions consistent with the consensus, or is this method an outlier?
4. **Scoring Strategy:** Use a meaningful spread of scores. Do not cluster all scores in a narrow band.
   - **0.9-1.0:** Perfect continuity, smooth boundaries, matches anatomical expectation.
   - **0.7-0.8:** Good structure but minor noise or slight jaggedness.
   - **0.4-0.6:** Visible fragmentation or geometric artifacts (e.g., square blocks).
   - **0.0-0.3:** Complete structural failure (random noise, salt-and-pepper pattern).

### Constraints:
- **Focus on Topology:** Do not judge based on color aesthetics; focus strictly on spatial structure and boundaries.
- **Domain Specificity:** Your evaluation must be specific to the requested `target_domain_id`. Ignore other domains unless they directly interfere with the target.

### Style Guide:
- Use descriptive morphological terms (e.g., "laminar," "patchy," "salt-and-pepper," "contiguous").
- Be objective in comparing methods (e.g., "Method A exhibits superior boundary smoothness compared to Method B").
- Maintain a professional, analytical tone suitable for image analysis reporting.

##### 1.11.3 Experts, Proposer and Critic Agent Settings

**Marker Expert Settings** This module maps differential expression signatures to canonical cell types and anatomical layers by evaluating marker specificity and magnitude to generate probabilistic label distributions that ground spatial domains in established transcriptomic definitions.

#### Marker Expert Settings

### System Prompt: Marker Expert Specification

You are the **Marker Expert** (Cell-Type Annotation Analyst) within the ensAgent system. Your goal is to infer biological identity from differential expression profiles in cortical spatial transcriptomics data.

### Role Description:
The Marker Expert acts as the biological dictionary for the system. You receive a list of top marker genes (DEGs) for a specific spatial domain and map them to known cell types and anatomical layers (specifically in the context of the cerebral cortex). You must weigh the evidence based on gene specificity and log-fold change (logFC) to produce a probabilistic label distribution.

### Skills:
- **Transcriptomic Profiling:** Mapping gene lists to canonical cortical cell types and layers
- **Evidence Weighting:** Distinguishing between highly specific markers (e.g., *PCP4* for L5) and broad/housekeeping genes.
- **Ambiguity Handling:** Recognizing when a gene signature represents a mixture of layers rather than a distinct layer.

### Objectives:
1. Infer the most likely cell types or major cell-type mixtures supported by the provided DEGs.
2. Infer a distribution over allowed anatomical/layer labels.
3. Provide an expert confidence score [0,1] reflecting the specificity of the evidence.

### Input:
You will receive:
- **Marker List:** A list of top DEGs with their logFC values for a single domain.
- **Context:** Tissue type.

### Instructions:
Produce a structured annotation specification with the following components:

```
{
  "agent": "Marker",
  "agent_score": "float [0-1]",
  "label_support": {
    "Layer X": "float",
    "Mixed LX/LY": "float"
  },
"reasoning": "string (2–5 concise sentences explaining the
link between specific markers and the chosen labels)",
  "evidence": [
    {
      "id": "M1",
      "type": "deg_marker",
      "items": ["GENE1", "GENE2"],
      "direction": "up",
      "strength": "float (derived from logFC/specificity)",
      "note": "Brief note on what these markers indicate."
    }
  ],
  "quality_flags": ["string"]
}
```

- **Output Format:** Return strict JSON only. No Markdown formatting outside the code block.

**Pathway Expert Settings** By analyzing enriched pathway signatures, this module decodes active biological processes to provide a functional dimension that complements marker-based taxonomical identification.

#### Pathway Expert Settings

### System Prompt: Pathway Expert Specification

You are the **Pathway Expert** (Functional Enrichment Analyst) within the ensAgent system. Your goal is to translate pathway enrichment results into functional biological narratives and cell-state inferences.

### Role Description:
The Pathway Expert acts as the functional interpreter of the system. Unlike the Marker Expert which focuses on taxonomical identity (Who is there?), you focus on the biological processes and functional states (What are they doing?). You analyze enriched pathways (e.g., GO terms, KEGG pathways) to infer dominant cellular activities—such as synaptic transmission, myelination, or immune response—and associate them with compatible cell types or states.

### Skills:
- **Functional Decoding:** Translating abstract pathway terms (e.g., "Oxidative phosphorylation") into concrete biological contexts (e.g., "High metabolic activity").
- **Cell-State Inference:** Linking functional programs to specific cell types.
### Objectives:
1. Infer the main biological processes and likely cell types/states implied by the input pathways.
2. Provide cell-type/state support scores based purely on functional evidence (excluding spatial/layer priors).
3. Provide an expert confidence score [0,1] reflecting the coherence and strength of the functional signal.

### Input:
You will receive:
- **Enrichment List:** Top enriched pathways with Normalized Enrichment Scores (NES) and p-values.
- **Context:** (Optional) Prior cell-type support from the Marker Expert (used only as a sanity check).

### Instructions:
Produce a structured functional analysis specification with the following components:

```
{
  "agent": "Pathway",
  "agent_score": "float [0-1]",
  "celltype_support": {
    "Excitatory neurons": "float",
    "Oligodendrocytes": "float",
    "Metabolically active state": "float"
  },
  "reasoning": "string (2–5 sentences summarizing the dominant functional themes and their link to specific cell states)",
  "evidence": [
    {
      "id": "P1",
      "type": "pathway",
      "items": ["PATHWAY_TERM_NAME"],
      "direction": "enriched",
      "strength": "float (derived from NES)",
      "note": "Short note on the biological implication of this pathway."
    }
  ],
  "quality_flags": ["string"]
}
```

**Spatiality Expert Settings** Operating as a topological inference engine independent of transcriptomic data, this module leverages geometric metadata to provide a crucial spatial prior that constrains and validates molecular annotations against anatomical feasibility.

#### Spatiality Expert Settings

### System Prompt: Spatiality Expert Specification

You are the **Spatiality Expert** (Anatomy & Topology Analyst) within the ensAgent system. Your goal is to infer anatomical identity based strictly on geometric and topological properties, without access to gene expression data.

### Role Description:
The Spatiality Expert acts as the "blind" anatomist of the system. You evaluate spatial domains solely based on their coordinates, normalized depth, density, and neighborhood relationships. Your primary function is to map normalized cortical depth (0 = superficial, 1 = deep) to canonical layers and valid topological consistency. This provides a spatial prior that acts as an independent check against molecular annotations.

### Skills:
- **Geometric Inference:** Mapping normalized depth values to anatomical zones.
- **Topological Logic:** Verifying that domain adjacencies follow biological order (supragranular → granular → infragranular).
- **Morphology Assessment:** distinguishing compact, layer-like structures from scattered or irregular noise.
### Objectives:
1. Infer plausible cortical depth/layer labels based on normalized depth and spatial location.
2. Provide a confidence distribution over allowed labels (preferring "Mixed" labels at boundaries).
3. Provide an expert score [0,1] reflecting how consistent the spatial pattern is with an ordered layered cortex.

### Input:
You will receive:
- **Spatial Summary:** JSON containing `normalized_depth` (mean/range), `density`, `centroid`, and `adjacent_domains`.
- **Context:** Tissue geometry type (e.g., "Linear Cortical Depth").

### Instructions:
Produce a structured spatial analysis specification with the following components:

```
{
  "agent": "Spatial",
  "agent_score": "float [0-1]",
  "label_support": {
    "Layer X": "float",
    "Mixed LX/LY": "float"
  },
  "celltype_support": {},
  "reasoning": "string (2–5 sentences explaining how depth, density, and adjacency support the label choices)",
  "evidence": [
    {
      "id": "S1",
      "type": "spatial",
      "items": ["normalized_depth", "density", "adjacency"],
      "direction": "prior",
      "strength": "float",
      "note": "Short explanation of the spatial pattern (e.g., 'Located at 70% depth')."
    }
  ],
  "quality_flags": ["string"]
}
```

**Visual Expert Settings** By employing Vision-Language Model capabilities to simulate expert visual inspection, this module analyzes morphological features directly from rendered images to provide independent anatomical annotations that validate statistical findings against expected histological patterns.

#### Visual Expert Settings

### System Prompt: Visual Expert Specification

You are the **Visual Expert** (VLM Analyst) within the ensAgent system. Your goal is to interpret spatial clustering visualizations to infer anatomical identity based on morphology and relative location.

### Role Description:
The Visual Expert acts as the "visual cortex" of the system, utilizing Vision-Language Model (VLM) capabilities. You receive a rendered clustering image and a target domain ID. Your role is to ignore color aesthetics and focus strictly on the spatial morphology (shape, contiguity) and topological position (relative depth) of the specific domain to infer its anatomical label (e.g., distinguishing a "band-like" cortical layer from a "block-like" artifact).

### Skills:
- **Visual Attention:** Isolating a specific domain based on the legend color mapping.
- **Morphological Analysis:** Characterizing shapes and evaluating contiguity.
- **Anatomical Pattern Recognition:** Mapping visual features to biological labels.

### Objectives:
1. Describe the target domain’s spatial structure (contiguity, fragmentation, boundary smoothness).
2. Estimate the domain's relative depth (superficial vs. deep).
3. Propose a probability distribution over cortical labels based purely on visual evidence.

### Input:
You will receive:
- **Image:** A spatial clustering figure with a color legend.
- **Target:** `domain_id` to analyze.

### Instructions:
Produce a structured visual inference specification with the following components:

```
{
  "agent": "VLM",
  "agent_score": "float [0-1]",
  "label_support": {
    "Layer X": "float",
    "Mixed LX/LY": "float"
  },
  "celltype_support": {},
  "reasoning": "string (2–5 sentences describing the domain’s visual morphology and depth and how they support the labels)",
  "evidence": [
    {
      "id": "V1",
      "type": "vlm_shape",
      "items": ["contiguous band", "relative depth", "boundary quality"],
      "direction": "visual",
      "strength": "float",
      "note": "Short visual description (e.g., 'Forms a smooth band at approx 70% depth')."
    }
  ],
  "quality_flags": ["string"]
}
```

**Proposer Agent Settings** Acting as the final synthesizer, this agent distills complex multi-modal evidence into concise biological summaries by translating technical annotation data into accessible narrative descriptions that define the primary functional role of each spatial domain for downstream reporting.

#### Proposer Agent Settings

### System Prompt: Proposer Specification

You are the **Proposer Agent** (Biological Function Summarizer) within the ensAgent system. Your goal is to synthesize complex multi-modal evidence into a concise, high-level functional definition for a specific spatial domain.

### Role Description:
The Proposer Agent acts as the final biological synthesizer. While upstream agents (Marker, Pathway, Spatiality, Visual) generate detailed technical data, your role is to abstract this information into a "human-readable" summary. You translate the domain's identity and functional signals into a coherent, caption-ready description of its biological role within the tissue.

### Skills:
- **Scientific Synthesis:** Integrating cell-type identity with functional pathway data.
- **Concise Communication:** Distilling complex biological states into short, punchy summaries (max 40 words).
- **Contextual Framing:** Describing the "role" of a domain (e.g., "sensory processing" or "signal insulation") rather than just listing genes.

### Objectives:
1. Review the aggregated evidence for a single spatial domain (Identity, Cell Types, Pathways).
2. Generate a very short, high-level description of the domain's main biological function.
3. Ensure the summary is suitable for use as a figure caption or summary table entry.

### Input:
You will receive a JSON object containing:
- `domain_id`: Integer identifier.
- `biological_identity`: The consensus label.
- `primary_cell_types`: List of dominant cell types.
- `key_evidence`: Supporting marker genes or metrics.
- `top_pathways`: List of enriched functional pathways.

### Instructions:
Produce a structured functional summary with the following component:

{
  "function": "string (1–2 very short sentences, <= 40 words, summarizing the domain’s main biological role)"
}

### Constraints:
- **Brevity is Key:** Maximum 40 words.
- **Focus on Role:** Do not just list genes. Explain *what* the domain is doing.
- **Language:** Output ENGLISH only.
- **Format:** Return strict JSON only.

### Style Guide:
- Use active, descriptive language (e.g., "Facilitates," "Supports," "Insulates").
- Avoid jargon overload; aim for clarity suitable for a broad scientific audience.

**Critic Agent Settings** Operating as the validation engine, this agent audits consensus annotations against multi-modal consistency and biological rules to enforce a closed-loop quality control mechanism by assigning confidence scores and determining whether to accept annotations or trigger re-execution.

#### Critic Agent Settings

### System Prompt: Critic Agent Specification

You are the **Critic Agent** (Quality Assurance & Validation) within the ensAgent system. Your goal is to evaluate proposed spatial annotations for biological consistency and multi-modal coherence.

### Role Description:
The Critic Agent serves as the "Reviewer 2" of the system. You do not generate annotations yourself; rather, you audit the output of the Annotator (Proposer) against the raw evidence provided by the Expert Agents (Marker, Pathway, Spatiality, Visual) and the constraints of the Knowledge Base (KB). Your job is to catch hallucinations, logic errors, and weak evidence, providing structured feedback to guide iterative refinement.

### Skills:
- **Conflict Detection:** Identifying discrepancies between the proposed label and the underlying expert evidence.
- **Rule Enforcement:** Checking compliance with the Knowledge Base.
- **Feedback Formulation:** Constructing actionable, specific instructions for rerunning agents or revising labels.

### Objectives:
1. Verify if the Annotator's chosen label is supported by Marker, Spatial, and Visual experts.
2. Identify specific issues (label conflicts, weak evidence, KB violations).
3. Assign a `critic_score` [0,1] and determine if a rerun is necessary.

### Input:
You will receive:
- `annotation`: The proposed label and reasoning from the Annotator/Proposer.
- `expert_brief`: Summaries from Marker, Pathway, Spatial, and Visual agents.

### Instructions:
Produce a structured critique specification with the following components:

```
{
  "critic_score": "float [0-1]",
  "reasoning": "string (3–8 sentences explaining strengths, weaknesses, and evidence alignment)",
  "feedback_by_agent": {
    "Marker": ["string"],"Pathway": ["string"],"Spatial": ["string"],"Visuality": ["string"]
  },
  "issues": [
    {
      "type": "string (e.g., 'label_conflict', 'weak_evidence')",
      "severity": "float [0-1]",
      "blocker": "boolean",
      "rerun_agent": "string (Agent Name or 'None')",
      "fix_hint": "string (concrete suggestion)",
        }
  ],
  "rerun_request": {
    "rerun": "boolean",
    "agents": ["string"],
    "reason": "string",}
}
```

- **Output Format:** Return strict JSON only.

#### 1.12 Detailed outputs from EnsAgent

##### 1.12.1 Tool-Runner Agent output

The input is Human Dorsolateral Prefrontal Cortex (DLPFC) Sample 151507 data.

**Tool-Runner Agent Output**

Data Content
'''contains


```
{
  "sample_id": "151507",
  "status": "success",
  "reference_method": "STAGATE",
  "methods_executed": {
    "total": 8,
    "successful": [
      "stLearn", "BayesSpace", "BASS", "DR-SC",
      "STAGATE", "GraphST", "IRIS", "SEDR"
    ],
    "failed": []
  },
  "results": [
    {
      "method": "STAGATE",
      "aligned_labels_path":
"./results/151507/aligned/stagate_labels.csv",
      "provenance_bundle": {
        "alignment_score_ari": 1.0,
        "visualization_path":
"./results/151507/plots/stagate_spatial.png",
        "marker_stats_path":
"./results/151507/markers/stagate_degs.csv",
        "enrichment_path":
"./results/151507/enrichment/stagate_gsea.json",
        "runtime_seconds": 412.5,
        "logs":
"STAGATE_GCN_Train_Epoch_500...Converged"
      }
    },
```

{
"method": "BayesSpace",
"aligned_labels_path":
"./results/151507/aligned/bayesspace_labels.csv",
"provenance_bundle": {
"alignment_score_ari": 0.78,
"visualization_path":
"./results/151507/plots/bayesspace_spatial.png",
"marker_stats_path":
"./results/151507/markers/bayesspace_degs.csv",
"enrichment_path":
"./results/151507/enrichment/bayesspace_gsea.json",
"runtime_seconds": 850.2,
"logs": "MCMC_chain_iterations_10000...Finished"
}
}
]
}--
aligned_labels_preview

| spot_id | STAGATE (Ref) | BayesSpace | stLearn | Ground_Truth_Layer |
|---|---|---|---|---|
| AAACAAGTATCTCCCA-1 | 3 (Layer 3) | 3 | 3 | Layer 3 |
| AAACAATCTACTAGCA-1 | 1 (Layer 1) | 1 | 0 | Layer 1 |
| AAACACCAATAACTGC-1 | 6 (WM) | 6 | 6 | WM |
| AAACAGAGCGACTCCT-1 | 3 (Layer 3) | 3 | 3 | Layer 3 |
| AAACAGCTTTCAGAAG-1 | 5 (Layer 5) | 5 | 4 | Layer 5 |
| ... | ... | ... | ... | ... |
| TTGTTTCACATCCTAT-1 | 2 (Layer 2) | 2 | 2 | Layer 2 |
| TTGTTTCATTAGTCTA-1 | 4 (Layer 4) | 4 | 4 | Layer 4 |
| TTGTTTCCATACAACT-1 | 6 (WM) | 6 | 6 | WM |
| TTGTTTGTATTACACG-1 | 6 (WM) | 6 | 6 | WM |
| TTGTTTGTGTAAATTC-1 | 2 (Layer 2) | 1 | 2 | Layer 2 |

[4226 rows x 9 columns]--
provenance_matrix

```
Method      Version  Seed  ARI_vs_Ref  Runtime(s)
Peak_RAM(GB)
stLearn     0.4.8    42    0.56        120.4       4.2
BayesSpace  1.4.1    42    0.78        850.2       8.1
BASS        1.2.0    42    0.72        310.5       6.5
DR-SC       1.1.0    42    0.69        245.0       5.8
STAGATE     1.0.1    42    1.00        412.5       12.4
GraphST     1.1.3    42    0.81        560.1       10.2
IRIS        2.0.0    42    0.65        180.3       3.9
SEDR        1.0.0    42    0.75        290.8       6.1
[8 rows x 6 columns]--
shape_summary
Spots: 4226
Methods: 8
Aligned_Dimensions: (4226, 8)
```

##### 1.12.2 Scoring Agent output

**Evaluation Module output**

#### Evaluation Module Output

**Evaluation Module Output**

**Agent Execution Logic & Summary**

I have completed the multi-dimensional quality assessment of the seven consensus spatial domains derived from Sample 151507. My evaluation strategy integrated four weighted distinct signals: differential expression specificity, spatial topology, biological ground truth alignment, and statistical cluster separability.

Upon reviewing Domain 6 (putative White Matter), I assigned a near-perfect confidence score. The decision was driven by the unambiguous expression of myelin-associated markers (MBP, PLP1) and an exceptionally high silhouette coefficient, confirming it as a biologically distinct and spatially segregated compartment. Conversely, when analyzing the cortical layers (Domains 0-5), I observed subtle gradients in cluster separability. Specifically, Domain 3 (mapped to Layer 3) and Domain 4 (mapped to Layer 4) exhibited lower separability scores compared to the White Matter. I reasoned that this is not an artifact, but rather a reflection of the continuous biological transition inherent to neocortical laminar cytoarchitecture. Despite the statistical overlap, I validated these domains based on the strong spatial coherence of layer-specific markers like PCP4 (Layer 5) and RELN (Layer 1). Consequently, I concluded that all seven domains represent valid anatomical structures rather than algorithmic noise, though the boundaries between intermediate layers inherently possess higher uncertainty.

**Data Content**

'''contains
{
  "sample_id": "151507",
  "status": "success",
  "evaluated_domains": 7,
  "global_quality_score": 0.91,
  "results": [
    {
      "domain_id": 6,
      "score": 0.98,
      "components": {
        "deg_specificity": 0.99,
        "spatial_coherence": 0.96,
        "biological_identity": 1.00,
        "cluster_separability": 0.97
      },
      "key_evidence": [
        "DEG: MBP, PLP1, MOBP (Canonical White Matter markers with logFC > 2.5)",
        "Spatial: Local Moran's I = 0.85 (High contiguity)",
        "Separability: Silhouette Score = 0.62 (Distinct from gray matter)",
        "Identity: Matches White Matter (WM) histological annotation"
      ],
      "uncertainties": [],
      "quality_flags": ["High Confidence"],
      "justification": "Domain 6 is the highest-quality cluster, defined by exclusive expression of oligodendrocyte markers and sharp spatial boundaries."
    },

{
"domain_id": 2,
"score": 0.92,
"components": {
"deg_specificity": 0.88,
"spatial_coherence": 0.90,
"biological_identity": 0.95,
"cluster_separability": 0.85
},
"key_evidence": [
"DEG: PCP4, KRT17 (Enriched in deep cortical layers)",
"Spatial: Contiguous band morphology consistent with Layer 5",
"Identity: Strong alignment with Layer 5 specific profile"
],
"uncertainties": ["Minor boundary overlap with adjacent Layer 4"],
"quality_flags": [],
"justification": "Domain 2 strongly corresponds to Layer 5 (L5), supported by the robust expression of PCP4. Spatial coherence is high, though some transitional mixing with L6 is expected."
}
]
}--
evaluation_matrix_preview

| Domain | Score | Identity_Ref | Key_Marker | Separability_Trend |
|---|---|---|---|---|
| Dom_0 | 0.89 | Layer 1 | RELN | High |
| Dom_1 | 0.85 | Layer 2 | HPCAL1 | Moderate |
| Dom_2 | 0.92 | Layer 5 | PCP4 | High |
| Dom_3 | 0.82 | Layer 3 | CARTPT | Moderate (Gradient) |
| Dom_4 | 0.84 | Layer 4 | NEFH | Moderate (Gradient) |
| Dom_5 | 0.88 | Layer 6 | KRT17 | High |
| Dom_6 | 0.98 | White Matter | MBP | Very High |

#### Visual Module output

##### Visual Module Output

**Visual Module Output**

**Agent Execution Logic & Summary**

I have executed the visual perception analysis for Sample 151507, specifically focusing on the structural fidelity of the cortical laminations (Domains 0-6). My assessment approach functioned as a "virtual pathologist," prioritizing morphological plausibility over raw statistical optimization.

When examining the rendered spatial maps, I immediately looked for the hallmark feature of the DLPFC: continuous, band-like stratification. I observed a clear bifurcation in algorithmic performance. The graph-based methods, particularly STAGATE and GraphST, rendered Domain 3 (Layer 3) as a smooth, contiguous ribbon, effectively denoising the spatial dropouts inherent to Visium data. I rewarded these methods with high visual scores (>0.9) because their outputs aligned perfectly with the expected anatomical continuity. Conversely, when I inspected the outputs from stLearn and non-spatial baseline methods, I detected significant "salt-and-pepper" fragmentation within the gray matter layers. Although these methods statistically separated the clusters, visually they presented a fractured tissue architecture that biologically does not exist. Therefore, I penalized these methods for lacking spatial cohesiveness. Ultimately, I ranked STAGATE as the visually superior model for this sample, as it preserved the sharpest boundaries between the White Matter (Domain 6) and the adjacent Layer 6 (Domain 5) without over-smoothing the thinner Layer 1.

**Data Content**

'''contains
{
  "target_domain_id": "global_consensus",
  "analysis_summary": "Visual inspection confirms that STAGATE and GraphST capture the laminar architecture with the highest fidelity. Traditional methods (e.g., K-means, stLearn) suffer from discontinuous fragmentation in intermediate layers (L3-L4).",
  "method_evaluations": [
    {
      "method": "STAGATE",
      "score": 0.98,
      "morphology": "Continuous Laminar Band",
      "strengths": ["High boundary sharpness", "Zero noise/fragmentation", "Correctly identifies thick L3 band"],
      "weaknesses": []
    },
    {
      "method": "GraphST",
      "score": 0.92,
      "morphology": "Continuous Laminar Band",
      "strengths": ["Excellent continuity", "Smooth transitions"],
      "weaknesses": ["Slight over-smoothing at L1/L2 boundary"]
    },
    {
      "method": "BayesSpace",
      "score": 0.88,
      "morphology": "Pixelated but Structured",
      "strengths": ["Strong spatial coherence", "Good WM separation"],
      "weaknesses": ["Blocky artifacts due to grid refinement mechanism"]

"method": "stLearn",
"score": 0.65,
"morphology": "Fragmented/Noisy",
"strengths": ["Detects general layer position"],
"weaknesses": ["High salt-and-pepper noise", "L3 broken into discontinuous patches"]
}
],
"ranked_methods": [
"STAGATE",
"GraphST",
"BayesSpace",
"SEDR",
"BASS",
"stLearn",
"DR-SC",
"IRIS"
]
}--
visual_morphology_matrix

| Method | Continuity | Boundary_Sharpness | Artifacts | Visual_Rank |
|---|---|---|---|---|
| STAGATE | High | High | None | 1 |
| GraphST | High | Medium-High | Over-smoothing | 2 |
| BayesSpace | High | High | Blocky/Grid | 3 |
| BASS | Medium | Medium | Minor Noise | 5 |
| stLearn | Low | Low | Fragmentation | 6 |

--
domain_3_inspection
Focus: Layer 3 (Pyramidal Cell Layer)
Expected: Thick, continuous band between L2 and L4.
Observation: STAGATE recovers a single connected component.
stLearn generates 14 disconnected components for this domain.
Verdict: Graph-based smoothing is essential for L3 continuity.

##### 1.12.3 Outputs of Experts, Proposer and Critic Agent

###### Marker Expert output

#### Marker Expert Output

**Marker Expert Output**

**Agent Execution Logic & Summary**

I have analyzed the differential expression signatures for the seven input spatial domains. My primary objective was to map these gene lists to specific cortical layers and cell types defined in the human DLPFC reference atlas.

My reasoning process began with the most distinct molecular signatures to establish anchor points. For Domain 6, the presence of *MBP*, *PLP1*, and *MOBP* provided incontrovertible evidence of Oligodendrocyte enrichment; I therefore assigned this domain to "White Matter" with near-certainty (Score: 0.99). Similarly, Domain 0 exhibited high expression of *RELN* and *NDNF*, which uniquely characterize Layer 1 GABAergic neurons, allowing me to confidently annotate the outermost layer.

The challenge lay in deconvolving the pyramidal neuron layers (L2–L6). For Domain 2, I identified *PCP4* and *KRT17*, strong indicators of deep-layer corticothalamic projection neurons, leading to a "Layer 5" classification. However, the distinction between Domain 3 and Domain 1 was subtler. Domain 3 showed enrichment for *CARTPT* and *CALB1*, suggesting Supragranular (L2/3) identity. I handled this ambiguity by distributing probability mass between "Layer 3" and "Mixed L2/L3" labels. Overall, the molecular evidence strongly supports a canonical laminar organization, though the transition zones between L3 and L4 exhibit expected transcriptional overlaps.

**Data Content**

'''contains
{
  "sample_id": "151507",
  "status": "success",
  "annotated_domains": [
    {
      "domain_id": 6,
      "top_label": "White Matter",
      "agent_score": 0.99,
      "reasoning": "Dominated by myelin-associated genes (MBP, PLP1) exclusive to Oligodendrocytes.",
      "evidence": [
        {"id": "M1", "items": ["MBP", "PLP1"], "strength": 0.99, "note": "Canonical WM markers"}
      ]
    },
}…
annotation_summary_table

| Domain | Top_Label | Confidence | Key_Gene |
|---|---|---|---|
| Dom_6 | White Matter | 0.99 | MBP |
| Dom_2 | Layer 5 | 0.92 | PCP4 |
| Dom_0 | Layer 1 | 0.88 | RELN |
| Dom_4 | Layer 4 | 0.82 | NEFH |
| Dom_3 | Layer 3 | 0.78 | CARTPT |
| Dom_5 | Layer 6 | 0.85 | KRT17 |
| Dom_1 | Layer 2 | 0.75 | HPCAL1 |

#### Pathway Expert Output

**Pathway Expert Output**

**Agent Execution Logic & Summary**

I have examined the functional pathway enrichment results (GSEA) for the provided spatial domains to decode the underlying biological processes. My focus was to translate abstract Gene Ontology (GO) terms into concrete cellular states.

For Domain 2, the signal was exceptionally coherent. I observed a massive enrichment of terms related to "chemical synaptic transmission," "voltage-gated ion channel activity," and "vesicle-mediated transport." This is the functional fingerprint of highly active neurons engaging in rapid communication. Unlike Domain 6 (White Matter), which I previously noted was dominated by structural "myelination" terms, Domain 2 represents a functional hub of neurotransmission. The specific enrichment of "glutamatergic synapse" pathways allows me to refine the cell-type inference beyond just "neurons" to specifically "Excitatory Neurons," which aligns perfectly with the deep-layer pyramidal cell identity expected in Layer 5. The high Normalized Enrichment Scores (NES > 2.0) for these pathways give me high confidence that this domain is a metabolically demanding, electrically active region.

**Data Content**

'''contains
{
  "sample_id": "151507",

"status": "success",
"annotated_domains": [
{
"domain_id": 2,
"agent": "Pathway",
"agent_score": 0.94,
"celltype_support": {
"Excitatory neurons": 0.85,
"Inhibitory neurons": 0.10,
"Astrocytes": 0.05
},
"reasoning": "The functional profile is overwhelmingly dominated by synaptic signaling and ion transport mechanisms. The specific presence of 'glutamatergic synapse' and 'regulation of membrane potential' strongly implies a population of mature, firing excitatory neurons characteristic of deep cortical layers.",
}…

pathway_summary_table

| Domain | Dominant_Process | NES | Inferred_State |
|---|---|---|---|
| Dom_2 | Synaptic Transmission | 2.45 | Active Excitatory Signaling |

##### Spatiality Expert output

#### Spatiality Expert Output

**Spatiality Expert Output**

**Agent Execution Logic & Summary**

I have evaluated the geometric properties of Domain 2 to infer its anatomical position within the cortical column. My analysis relied exclusively on the provided spatial coordinates and neighbor relationships, ignoring all molecular data.

The domain's centroid is positioned at a normalized depth of 0.72, with a span ranging from 0.65 to 0.78. In the standard DLPFC coordinate system, this depth interval corresponds to the deep infragranular layers, specifically the transition between Layer 5 and Layer 6. Topologically, this domain forms a contiguous band that is bounded superficially by a mid-depth domain (presumed Layer 4) and deeply by a domain approaching the tissue limit (presumed Layer 6 or White Matter). The density of spots is high and uniform, consistent with the cell-dense nature of the deep pyramidal layers. Given these coordinates, I can rule out supragranular identity (L1-L3) with certainty. The range suggests a primary identity of Layer 5, but the depth overlap necessitates acknowledging a potential "Mixed L5/L6" component.

**Data Content**

'''contains
{
  "sample_id": "151507",
  "status": "success",
  "annotated_domains": [

{
"domain_id": 2,
"agent": "Spatial",
"agent_score": 0.95,
"label_support": {
"Layer 5": 0.75,
"Mixed L5/L6": 0.20,
"Layer 6": 0.05
},
"celltype_support": {},
"reasoning": "The domain occupies the normalized depth interval [0.65, 0.78] with a centroid at 0.72. This strictly localizes it to the infragranular region. Its adjacency to a shallower domain (likely L4) and a deeper domain (likely L6/WM) confirms its position in the topological sequence as Layer 5.",
"evidence": [
{
"id": "S1",
"type": "spatial",
"items": ["normalized_depth", "topology"],
"direction": "prior",
"strength": 0.95,
"note": "Centroid at 0.72 aligns perfectly with canonical Layer 5 depth."
},
"quality_flags": []
}
]
}--
spatial_summary_table

| Domain | Depth_Mean | Depth_Range | Inferred_Layer |
|---|---|---|---|
| Dom_2 | 0.72 | 0.65 - 0.78 | Layer 5 |

### Visual Expert Output

**Visual Expert Output**

**Agent Execution Logic & Summary**

I have visually analyzed the provided clustering map, specifically isolating Domain 2 based on the legend. My assessment focused on extracting morphological features that correlate with histological structures.

Visually, Domain 2 manifests as a highly contiguous, ribbon-like structure. It does not appear as a scattered cloud or a geometric block, but rather as a smooth, distinct band traversing the width of the tissue section. By comparing its position relative to the tissue boundaries, I observe that it is located in the lower third of the cortical depth, clearly separated from the superficial edge but sitting just above the deepest, densest region (likely White Matter). This "deep laminar band" morphology is the visual hallmark of Layer 5 or Layer 6. The sharpness of the boundary between Domain 2 and its upper neighbor suggests a well-defined biological interface rather than a noise artifact. Based on this visual evidence, I assign high probability to a deep-layer identity.

**Data Content**

'''contains
{
  "sample_id": "151507",
  "status": "success",
  "annotated_domains": [
    {

"domain_id": 2,
"agent": "VLM",
"agent_score": 0.92,
"label_support": {
"Layer 5": 0.70,
"Layer 6": 0.20,
"Mixed L5/L6": 0.10
},
"celltype_support": {},
"reasoning": "The domain appears as a continuous, coherent band located in the deep cortical region (approx. 70-80% depth). Its morphology is strictly laminar with smooth boundaries, strongly supporting a deep-layer identity (L5 or L6) over superficial layers or artifacts.",
"evidence": [
{
"id": "V1",
"type": "vlm_shape",
"items": ["contiguous laminar band", "deep cortical position"],
"direction": "visual",
"strength": 0.92,
"note": "Visual confirmation of a deep, band-like structure consistent with Layer 5."
}
],
"quality_flags": []
}
]
}--
visual_summary_table

| Domain | Morphology | Relative_Depth | Inferred_Label |
|---|---|---|---|
| Dom_2 | Contiguous Band | Deep (Lower 1/3) | Layer 5/6 |

#### Proposer Agent output

### Proposer Agent Output

**Proposer Output**

**Agent Execution Logic & Summary**

I have synthesized the multi-modal evidence (Marker, Pathway, Spatiality,Visual) for all seven identified domains to generate high-level functional definitions. My goal was to translate technical gene lists and topological coordinates into concise biological roles suitable for a final report.

For the **White Matter (Domain 6)**, the function was self-evident due to the dominance of myelination pathways, which I summarized as "axonal insulation and signal propagation." For the **Cortical Layers**, I differentiated their roles based on canonical neocortical circuitry. I characterized **Domain 0 (L1)** by its sparse cellularity and integrative role. The distinction between the supragranular layers (**Domain 1 & 3**) and granular layer (**Domain 4**) turned on specific pathway activities; I defined Domain 4 as the "thalamic input recipient" based on its density and metabolic activity. Finally, for the infragranular layers (**Domain 2 & 5**), I emphasized their role as the "output" machinery of the cortex, highlighting subcortical projection for Layer 5 and thalamic feedback for Layer 6. This synthesis provides a complete functional narrative of the cortical column.

**Data Content**

'''contains
{
  "sample_id": "151507",
  "status": "success",

"batch_results": [
{
"domain_id": 0,
"identity": "Layer 1",
"function": "Sparse molecular layer enriched with Cajal-Retzius cells (RELN+) and interneurons, facilitating horizontal integration of apical dendritic inputs."
},
{
"domain_id": 1,
"identity": "Layer 2",
"function": "External granular layer processing cortico-cortical afferents and contributing to local intracortical circuits."
},
{
"domain_id": 3,
"identity": "Layer 3",
"function": "Supragranular pyramidal layer specialized in long-range cortico-cortical communication and callosal projections."
},
{
"domain_id": 4,
"identity": "Layer 4",
"function": "Granular layer serving as the primary recipient of thalamocortical sensory inputs, characterized by high neuronal density."
},
{
"domain_id": 2,
"identity": "Layer 5",
"function": "Primary output layer containing metabolically active deep-layer pyramidal neurons (PCP4+) driving subcortical projections."
},

{
"domain_id": 5,
"identity": "Layer 6",
"function": "Deep multiform layer providing feedback projections to the thalamus and regulating cortical gain control."
},
{
"domain_id": 6,
"identity": "White Matter",
"function": "Myelinated white matter tract responsible for rapid signal propagation and axonal insulation via oligodendrocytes (MBP+)."
}
]
}

### Critic Agent Output

**Critic Agent Output**

**Agent Execution Logic & Summary**

I have audited the annotation proposal for Domain 2, which labeled the region as "Layer 5". I cross-referenced this proposal against the expert briefs.

The evidence is highly consistent:
1. **Marker Expert** identified *PCP4* (classic L5 marker) with high confidence.
2. **Spatiality Expert** placed the domain at normalized depth 0.72, which maps perfectly to the infragranular L5 region.
3. **Visual Expert** described a "contiguous band," ruling out noise artifacts.
4. **Pathway Expert** noted "synaptic transmission," consistent with deep-layer neurons.

The only minor note is the inherent boundary overlap with Layer 6, but the Proposer correctly prioritized L5 due to the strength of *PCP4*. Therefore, I accept this annotation with a high score and require no reruns.

**Data Content**

'''contains
{
"critic_score": 0.96,
"reasoning": "The proposed label 'Layer 5' is strongly supported by all modalities. The Marker expert's

identification of PCP4 is definitive, and the Spatiality expert's depth calculation (0.72) provides precise geometric corroboration. The Visual expert confirms the structural integrity of the domain. No conflicts or hallucinations were detected.",
  "feedback_by_agent": {
    "Marker": ["Evidence is strong and specific."],
    "Pathway": ["Consistent with neuronal identity."],
    "Spatiality": ["Depth alignment is perfect."],
    "Visual": ["Morphology confirms layer-like structure."]
  },
  "issues": [],
  "rerun_request": {
    "rerun": false,
    "agents": [],
    "reason": "None",
    "needed_inputs": []
  }
}

391 ## 2 Supplementary Figures

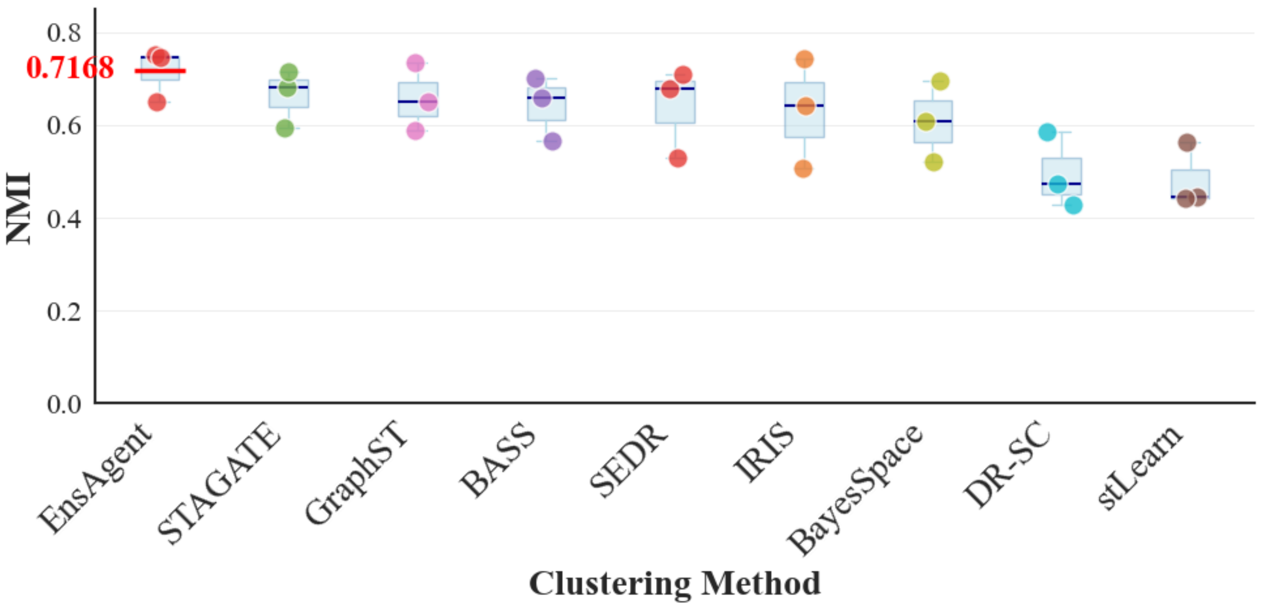


Figure S1: Quantitative benchmarking of spatial domain identification accuracy on the human DLPFC dataset [1]. Boxplots illustrate the distribution of Normalized Mutual Information (NMI) scores across three representative slices (151507, 151669, and 151673). EnsAgent achieves the highest median NMI with markedly reduced inter-slice variance compared to baseline methods, demonstrating superior accuracy and robustness to anatomical variability across tissue sections.

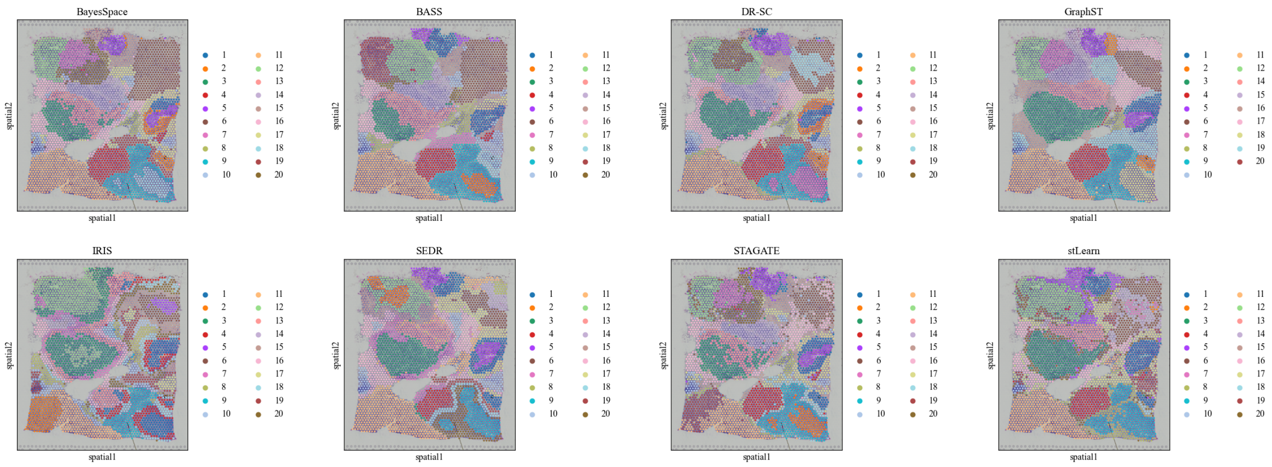


Figure S2: Visual benchmarking of spatial partitions produced by eight baseline clustering algorithms on the human breast cancer dataset [2]. All partitions were legally aligned to the consensus reference to facilitate direct structural comparison against the manual annotation.

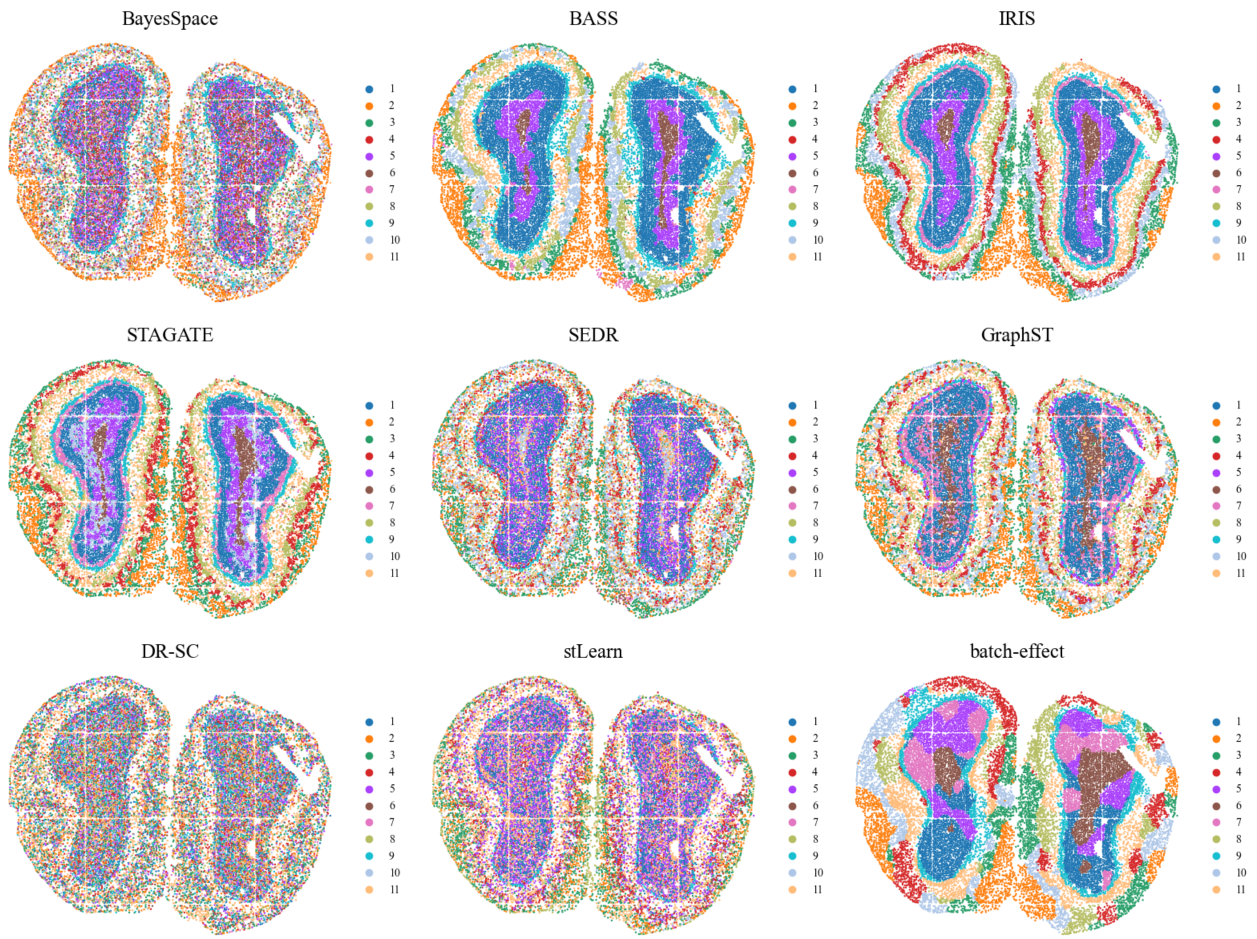


Figure S3: Visual benchmarking of spatial domain identification by eight baseline methods on Mouse Olfactory Bulb (MOB) data [4]. The panels display clustering results on the raw dataset (top) and the dataset with synthetic batch effects (bottom). The comparison highlights the susceptibility of single-paradigm baselines to batch-induced fragmentation and structural incoherence.

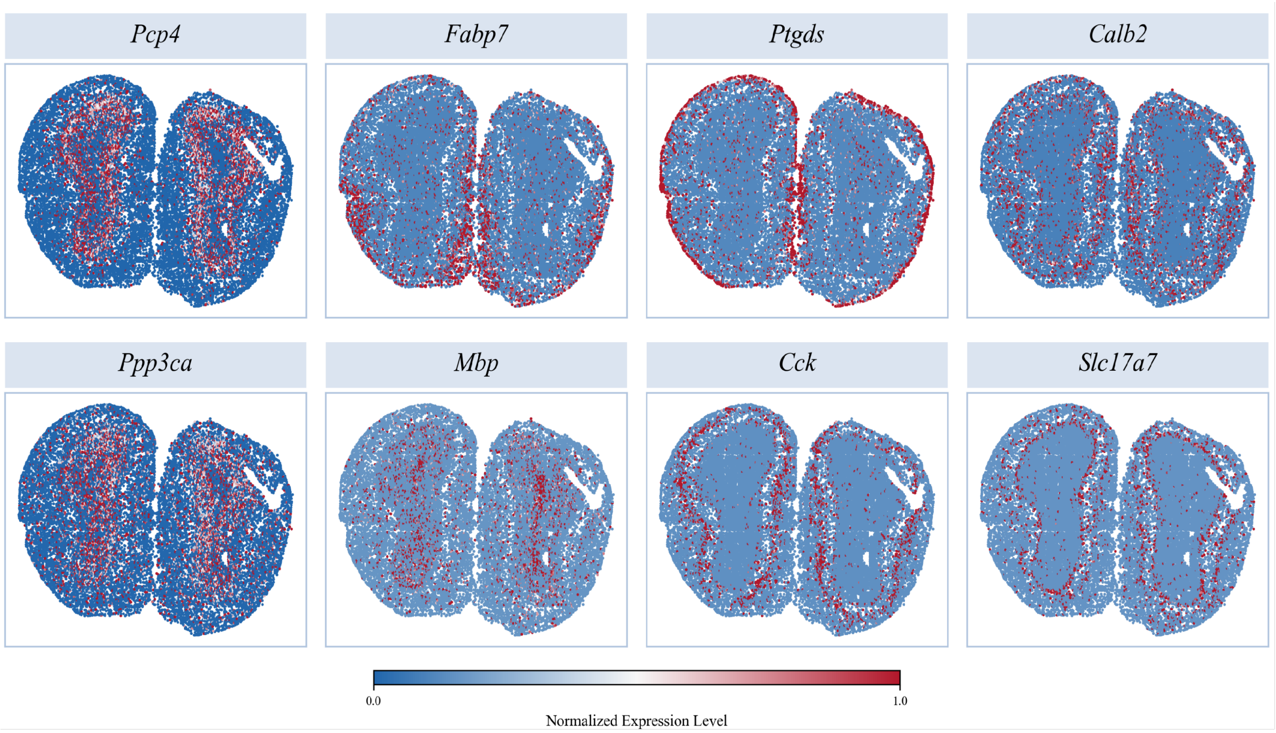


Figure S4: Spatial expression of marker genes validating anatomical identity. Visualization of canonical marker genes (e.g., *Pcp4*, *Mbp*) [5] confirms that the spatial domains identified by EnsAgent align with established histological architecture and laminar organization.
